# Massively parallel kinetic profiling of natural and engineered CRISPR nucleases

**DOI:** 10.1101/696393

**Authors:** Stephen K. Jones, John A. Hawkins, Nicole V. Johnson, Cheulhee Jung, Kuang Hu, James R. Rybarski, Janice S. Chen, Jennifer A. Doudna, William H. Press, Ilya J. Finkelstein

## Abstract

Engineered *Streptococcus pyogenes* (*Sp*) Cas9s and *Acidaminococcus sp*. (*As*) Cas12a (formerly Cpf1) improve cleavage specificity in human cells. However, the fidelity, enzymatic mechanisms, and cleavage products of emerging CRISPR nucleases have not been profiled systematically across partially mispaired off-target DNA sequences. Here, we describe NucleaSeq— <u>nuclea</u>se digestion and deep <u>seq</u>uencing—a massively parallel platform that measures cleavage kinetics and captures the time-resolved identities of cleaved products for more than ten thousand DNA targets that include mismatches, insertions, and deletions relative to the guide RNA. The binding specificity of each enzyme is measured on the same DNA library via the chip-hybridized association mapping platform (CHAMP). Using this integrated cleavage and binding platform, we profile four *Sp*Cas9 variants and *As*Cas12a. Engineered Cas9s retain wtCas9-like off-target binding but increase cleavage specificity; Cas9-HF1 shows the most dramatic increase in cleavage specificity. Surprisingly, wtCas12a—reported as a more specific nuclease in cells—has cleavage specificity similar to wtCas9 *in vitro*. Initial cleavage position and subsequent end-trimming vary across nucleases, guide RNA sequences, and position and base identity of mispairs in target DNAs. Using these large datasets, we develop a biophysical model that reveals mechanistic insights into off-target cleavage activities by these nucleases. More broadly, NucleaSeq enables rapid, quantitative, and systematic comparison of the specificities and cleavage products of engineered and natural nucleases.

## Introduction

RNA-guided CRISPR-associated (Cas) nucleases have ushered in the gene editing revolution. *S. pyogenes* Cas9 —the most widely used Cas nuclease—is guided to a target DNA via a ∼100-nt single guide RNA (sgRNA) (Hsu et al., 2013; Jinek et al., 2012). The first 20 nucleotides (nts) of the 5’ end of the sgRNA are complementary to the target DNA, referred to as the “protospacer” sequence. Cas9 interrogates potential genomic sites by first recognizing a three-nucleotide NGG protospacer adjacent motif (PAM), followed by propagation of an R-loop from the PAM, and finally activation of the nuclease domains (Gong et al., 2018; Jiang et al., 2016; Jinek et al., 2012; Sternberg et al., 2015). Cas9 also binds and cleaves off-target sites that are partially complementary to the sgRNA, resulting in unanticipated mutations, large-scale genomic deletions, and gross chromosomal rearrangements (Anderson et al., 2018; Cullot et al., 2019; Fu et al., 2013). Intense efforts have been focused on mitigating these off-target activities to improve gene editing specificity.

Engineered and natural Cas9 variants, as well as new subtypes of Cas nucleases, have been reported to reduce off-target DNA cleavage in cells (Amrani et al., 2018; Chen et al., 2017; Edraki et al., 2018; Kleinstiver et al., 2016a; Lee et al., 2018; Ran et al., 2015; Shmakov et al., 2015; Slaymaker et al., 2016; Smargon et al., 2017; Wu et al., 2018; Zetsche et al., 2015). For example, a series of engineered *Sp*Cas9s improve cleavage specificity by reducing protein interactions with either DNA strand or by modulating activation of the nuclease domains (Chen et al., 2017; Kleinstiver et al., 2016a; Slaymaker et al., 2016). In addition, the recently-discovered Cas12a is reported to be more specific than *Sp*Cas9 at some genomic sites in human cells (Kim et al., 2016; Zetsche et al., 2015). However, no single experimental strategy exists to directly benchmark nucleases by systematically measuring the key enzyme-intrinsic binding and cleavage specificities across partially matched DNA targets. Instead, nuclease specificity is typically assessed in cells by the extent of indel formation at the programmed target and a few known or putative off-target genomic sequences. Such approaches provide a broad overview of off-target nuclease activity but are limited by several critical shortcomings. First, these approaches do not provide a systematic comparison of how the mispaired base, its position within the R-loop, and the type of mispair—a mismatch, insertion, or deletion—impacts nuclease binding, cleavage rate, and nucleolytic DNA trimming. Second, the off-target binding specificity of high-fidelity nucleases has not been measured comprehensively. Third, these approaches do not differentiate enzyme-intrinsic kinetic parameters from confounding factors such as the nuclease delivery method and exposure time, genetic context, cell cycle phase, and dominant DNA repair pathways. Next-generation sequencing (NGS)-based *in vitro* approaches can address some of these limitations, but existing methods rely on sequence-limited genomic DNA substrates, generally capture read counts rather than cleavage rates, and do not report on the resulting DNA ends or their processing (Crosetto et al., 2013; Guenther et al., 2013; Kim et al., 2015; Tsai et al., 2015, 2017). We set out to develop an unbiased, massively-parallel, and general strategy to measure both DNA binding and cleavage specificity for benchmarking next-generation CRISPR-Cas nucleases.

Here, we describe NucleaSeq—<u>nuclea</u>se digestion and deep <u>seq</u>uencing—as a rapid and massively parallel platform for measuring cleavage kinetics by CRISPR-Cas nucleases. NucleaSeq also captures the time-resolved identities of cleaved products for a large library of fully and partially sgRNA-matched DNAs. The same DNA sequence library is used to measure the binding specificity of each enzyme via the chip-hybridized association mapping platform (CHAMP). Coupling NucleaSeq and CHAMP, we measured the cleavage and binding specificities of four *Sp*Cas9 variants and *Acidaminococcus sp*. Cas12a (wtCas12a) for DNAs containing mismatches, insertions, and deletions. Engineered Cas9s, most notably Cas9-HF1, dramatically increase cleavage specificity but provide minimal improvement to overall binding specificity. Surprisingly, wtCas12a—reported as a more specific nuclease in cells—resembles the cleavage specificity of wtCas9 *in vitro*. This suggests that wtCas12a’s reduced off-target cleavage in cells may stem from its overall slower cleavage rate (10-to 40-fold slower than wtCas9) or other cell-specific factors (i.e., expression levels, chromatin and cell cycle state, etc.). Both Cas9 and Cas12a produce variable ssDNA overhangs. Initial cleavage position and subsequent end-trimming vary with the nuclease, sgRNA sequence, and the mispairs between the sgRNA and the target DNA. By programming mispairs between sgRNA and target DNA, these nucleases can generate incompatible DNA ends without slowing cleavage, ultimately biasing cellular repair outcomes. Using these large datasets, we develop a biophysical model that gives additional mechanistic insights into off-target cleavage activities and provides a quantitative framework for comparing emerging CRISPR nucleases. More broadly, NucleaSeq and CHAMP enable rapid, quantitative, and systematic comparison of the specificities and cleavage products of engineered and natural nucleases.

## Results

### A massively-parallel platform for measuring nuclease cleavage and binding specificities

We set out to systematically evaluate the DNA cleavage and binding specificities of five CRISPR-Cas nucleases: wild-type *S. pyogenes* Cas9 (wt), three engineered *Sp*Cas9 variants (enhanced eSp1.1, high fidelity HF1, hyper accurate Hypa), and *Acidaminococcus species* Cas12a (wt, also known as Cpf1) (Figure 1A and S1A) (Chen et al., 2017; Jinek et al., 2012; Kleinstiver et al., 2016a; Slaymaker et al., 2016; Zetsche et al., 2015). For NucleaSeq, we synthesized a pooled library that contains >10^4^ barcoded DNAs that randomize the PAM or have up to two edits (i.e., mismatches, insertions or deletions) relative to the sgRNA (Figure 1B, 1C, Supplemental File 1). Each target is flanked by error-correcting barcodes that uniquely identify the two DNA fragments after nuclease-catalyzed cleavage (Hawkins et al., 2018). The PCR-amplified library is incubated with a ten-fold excess of each CRISPR holoenzyme (e.g., Cas9 or Cas12a ribonucleoprotein) (Figure S1B and S1C). This high molar excess ensures that we measure single-turnover kinetics. The reaction is sampled as a function of time, quenched in a stop solution, and deproteinized to release the cut DNA products (Figure 1D and S1D). Each time point is prepared for Illumina-based next-generation DNA sequencing (NGS); adapter ligation includes gap-filling 5’ DNA overhangs and trimming 3’ overhangs. The adapter also includes an outer barcode to identify each time point prior to sequencing via the standard Illumina workflow. The resulting datasets are analyzed via the NucleaSeq bioinformatics pipeline.

**Figure 1:**
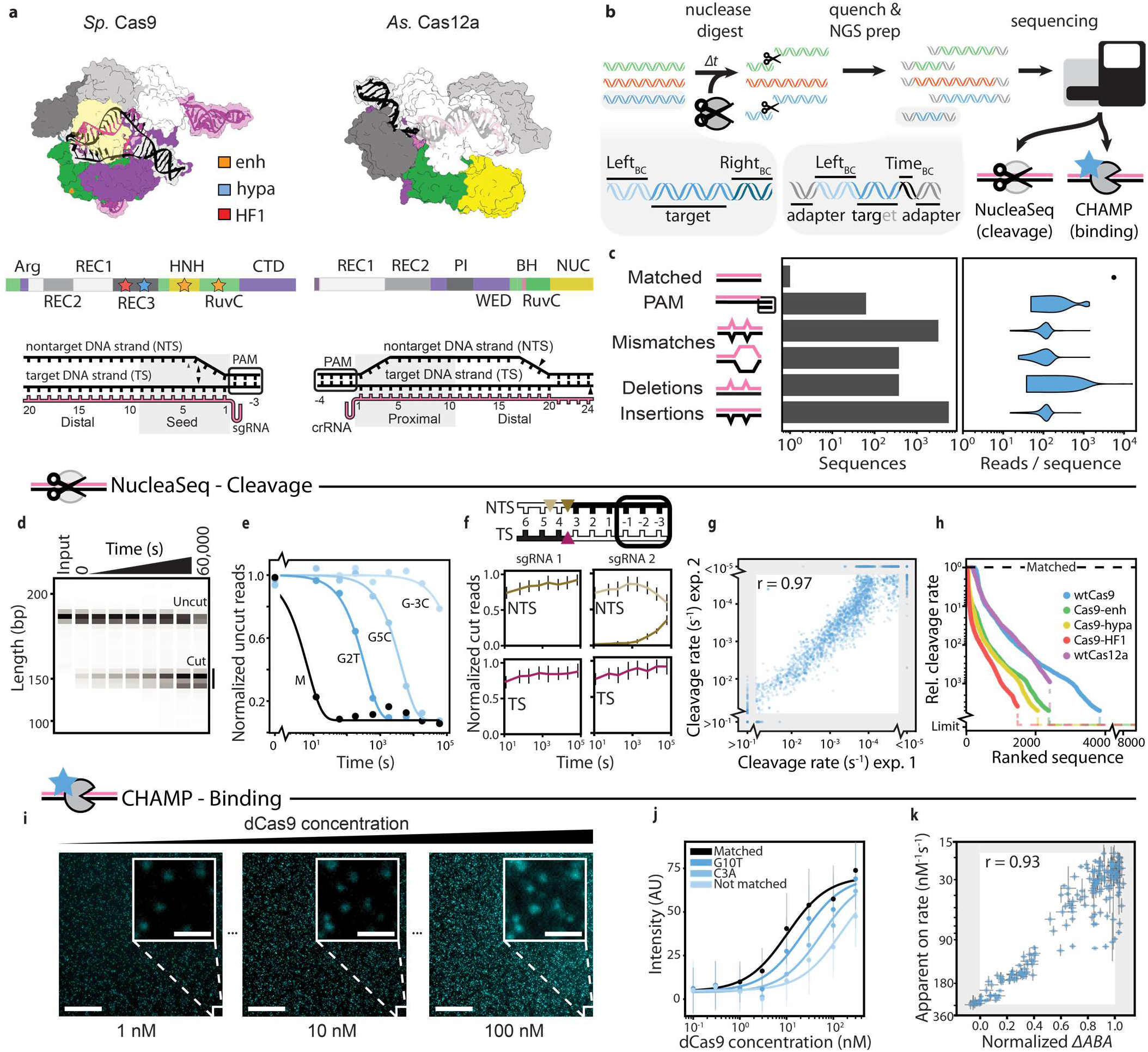
Overview of the integrated NucleaSeq and CHAMP platform. **(a)** Crystal structures and domain maps of Cas9 and Cas12a ribonucleoprotein complexes (RNPs; PDB: 5F9R, 5B43). Stars indicate clusters of mutations for engineered Cas9 variants. Triangles in the R-loops indicate reported cleavage sites. **(b)** For NucleaSeq, a synthetic library of partially mispaired DNA targets is digested with a CRISPR-Cas nuclease under single-turnover conditions. Each DNA incorporates unique left and right barcodes. An outer barcode is added to each timepoint prior to next-generation DNA sequencing (NGS). The NGS chips are recovered for profiling DNA-binding specificity via CHAMP. **(c)** Each DNA library includes a randomized PAM and up to two edits (i.e., mismatches, insertions, or deletions) relative to the corresponding gRNA. Right: distribution of reads for each type of mispaired target. Two DNA library-gRNA pairs were analyzed for each RNP. **(d)** The time course of a wtCas9 nuclease reaction (sgRNA 1) is resolved by capillary electrophoresis. **(e)** The cleavage rate is computed by fitting the time-dependent depletion of uncut library members (circles) to a single exponential (solid line). **(f)** The time-dependent distribution of Cas9-generated cut sites in the target (TS) and nontarget strands (NTS) of a matched target is reported by the cut fragments (black in diagram). With sgRNA 1, wtCas9 produces a blunt overhang between the 3^rd^ and 4^th^ nucleotides (left). With sgRNA 2, wtCas9 initially produces a one-nucleotide 5’ overhang that is trimmed towards a blunt cut (right). Line colors correspond to the indicated cut positions (triangles in diagram). Error bars: SEM based on 150 library members containing the matched DNA sequence. **(g)** Cleavage rates are highly reproducible for all experiments (wtCas9 data shown, r = Pearson correlation coefficient). Gray regions contain sequences that exceed the dynamic range of the experiment and are therefore excluded from the correlation. **(h)** The relative cleavage rate of all mismatched targets (matched target cleavage rate divided by the corresponding rate) for all five nucleases. Limit = relative cleavage rate that is beyond the limit of detection. **(i)** CHAMP reports the apparent binding affinity of nuclease-inactive CRISPR enzymes. DNA libraries attached to the surface of a sequenced NGS chip are incubated with increasing concentrations of a fluorescent dCas9 (cyan puncta). Scale bar: 50 μm, inset: 5 μm. The sequence of each DNA cluster is computationally identified by comparison to the NGS output. **(j)** The mean fluorescence intensity of DNA clusters with the same DNA sequence (symbols) are fit to a Hill equation (lines) to determine the apparent binding affinity (ABA, dCas9 shown). AU: arbitrary fluorescence units. Error bars: SD of DNA clusters containing the same DNA sequence. **(k)** The ΔABAs measured via CHAMP are highly correlated with the dCas9 on-rates as reported via a related high-throughput assay (Boyle et al., 2017). ΔABAs are change in apparent binding affinity from a matched DNA normalized to that of a scrambled DNA. X-axis error bars: SD as measured by bootstrap analysis (ΔABA); Y-axis: SE of the mean.

The bioinformatics pipeline first identifies cut and uncut (full-length) library members by their flanking barcode(s). The read counts for each library member are normalized across time points and between replicates by comparing to read counts of ∼150 negative control DNA sequences that are not recognized by any of the nucleases. Since Cas9 and Cas12a cleave DNA at a constant rate under single turnover conditions, substrate depletion follows a single exponential decay (Sternberg et al., 2014; Strohkendl et al., 2018). We determine the cleavage rate for each library member by fitting the time-dependent depletion of uncut DNAs to a single exponential function (Figure 1E) (Guenther et al., 2013). As expected, all nucleases cleave their matched DNA substrate rapidly (*k_c_* ≥0.1 s^−1^ for wtCas9, Figure 1E), but even a single mismatch between the target DNA and the sgRNA can reduce cleavage rates by >10^4^-fold (i.e., a G_5_ to C_5_ substitution, G5C, for wtCas9 in Figure 1E). The two cut products produced from the cleavage of each library member are then compared to identify the cut site and time-dependent trimming of the DNA ends (Figure 1F). The cleavage rates and cut product distributions were highly reproducible across two NucleaSeq biological replicates for both Cas9 and Cas12a nucleases (Figure 1G and S1E). The cleavage specificity—a ratio of the cleavage rate of a mismatched target to that of the matched target—is a simple way to benchmark nucleases. A large cleavage specificity indicates that the mispaired target is cut significantly slower than the matched target under saturating enzyme concentrations. Figure 1H summarizes the cleavage specificity of the five nucleases tested here. All engineered Cas9 variants outperform wtCas9 with Cas9-HF1 showing the highest specificity for all mismatched targets. Surprisingly, wtCas12a retains similar cleavage specificity to wtCas9. The cleavage rates are then compared to off-target DNA binding affinity for the same DNA libraries using the massively-parallel Chip-Hybridized Association Mapping Platform (CHAMP) (Jung et al., 2017).

CHAMP is used to measure the apparent binding affinity of CRISPR-Cas nucleases to DNA clusters on the surface of regenerated NGS chips (Figure 1). Following NGS, all surface-tethered DNA clusters are regenerated via a single round of primer extension (Jung et al., 2017). A subset of the clusters is fluorescently labeled for computational alignment of fluorescent images to the NGS sequencing information. The sequence identity of each DNA cluster is thus decoded by the CHAMP analysis pipeline. To measure off-target DNA binding, a fluorescent nuclease-dead Cas protein (e.g. dCas9 in Figure 1K) is incubated in the flowcell at increasing protein concentrations. The corresponding increase in fluorescent signal at each DNA cluster is fit to a hyperbolic function to reveal an apparent binding affinity (ABA) for the protein to each underlying DNA sequence (Figure 1J). The ABAs are highly reproducible across two biological CHAMP replicates (Figure S1F). Moreover, CHAMP-derived ABAs are strongly correlated with a prior report of dCas9 on-rates for mismatched DNA sequences (Figure 1K) (Boyle et al., 2017). We conclude that ABAs—and the underlying DNA binding specificity of Cas9—is largely driven by differences in the on-rates for different DNA sequences. More broadly, NucleaSeq and CHAMP use the same library of DNA sequences. Thus, coupled cleavage and binding information identify sequence-specific mechanisms of nuclease fidelity. A detailed comparison of the cleavage and binding specificities of these five nucleases is described below.

### Systematic analysis of wtCas9 binding and cleavage specificity

We first focused on the cleavage and binding specificity of wtCas9 (Figure 2 and S2). wtCas9 was loaded with one of two sgRNAs (Table S1), and then incubated with the corresponding DNA library for over 16 hours. Individual time points were sequenced to obtain the cleavage rates that spanned the entire detectable range of *k_c_* ∼10^−1^ to ∼10^−5^ s^−1^. The cleavage rate for the matched DNA (≥0.1 s^−1^) agrees closely with a previously-reported overall kinetic rate constant of 0.2 s^−1^, which is limited by the rate of R-loop propagation (Gong et al., 2018). To measure off-target DNA binding, increasing concentrations of wt dCas9 were incubated in regenerated MiSeq chips harboring the sequenced DNA library. We detected no DNA binding at the lowest dCas9 concentration (100 pM), while the DNA clusters appeared completely saturated at the highest dCas9 concentration (300 nM). Apparent binding affinities (ABAs) were highest for the matched DNA sequence and lowest for scrambled control DNAs that were not complementary to the sgRNA or lacked the NGG protospacer adjacent motif (PAM, see below). Consistent with prior reports *in vitro* and *in vivo*, dCas9 had a high apparent binding affinity for partially mismatched target DNA sequences. Our results were strongly correlated between two biological replicates and with the binding affinities measured via another high-throughput method (r=0.93, Figure 1G and S1E) (Boyle et al, 2017). Overall, we observed that Cas9 bound 70% of the library with a higher affinity than nonspecific DNA but could only cleave 60% of the same DNA library, indicating that cleavage is only partially determined by DNA binding.

**Figure 2:**
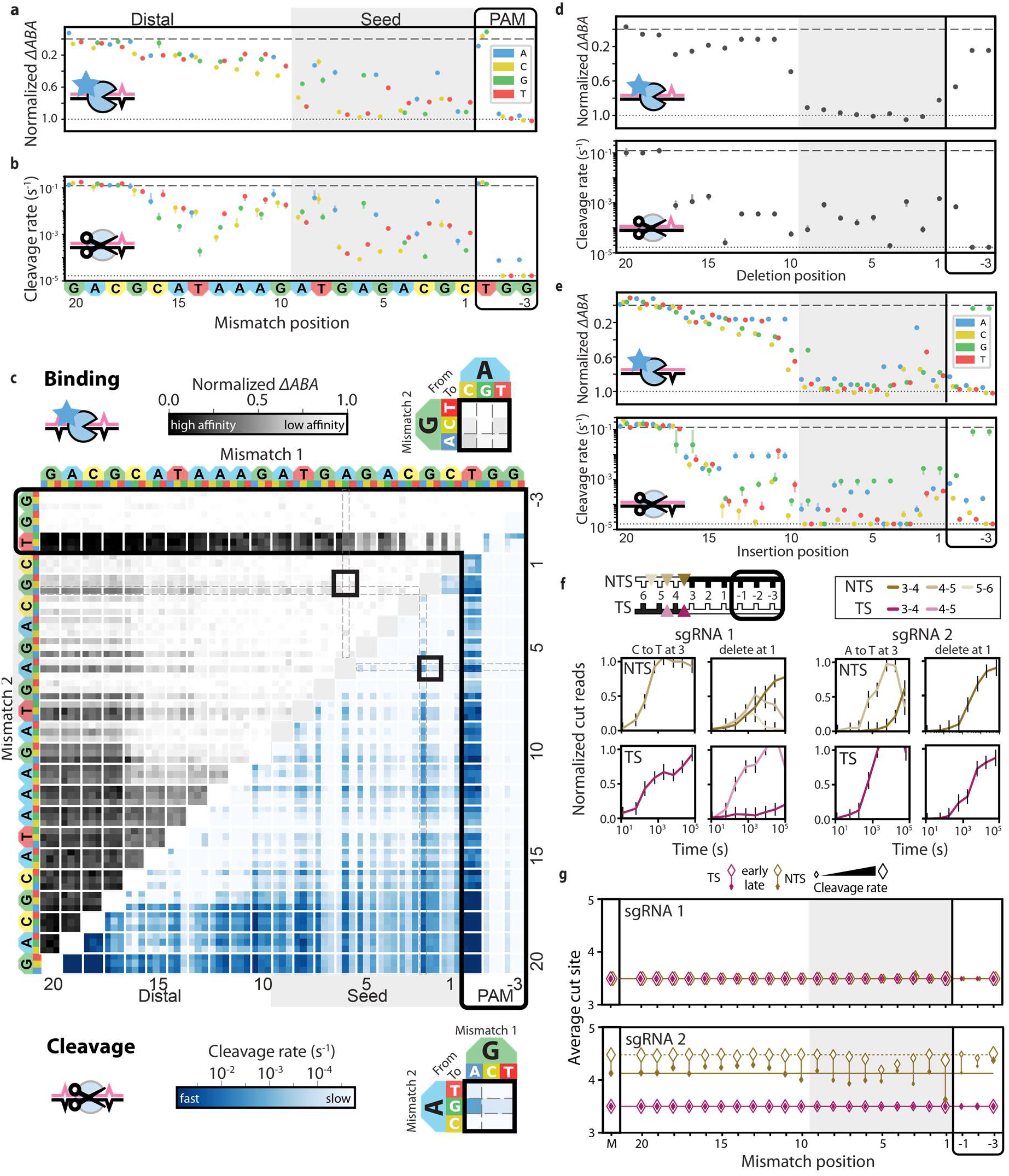
Comprehensive analysis of off-target wtCas9 DNA binding and cleavage. **(a)** dCas9 ΔABAs for all DNAs with a single mismatch relative to sgRNA 1. Dotted line: normalized matched target ΔABA (0); dashed line: scrambled DNA ΔABA (negative control, 1). Error bars: SD as measured by bootstrap analysis. **(b)** Cas9 cleavage rates for the same targets as in (a). Dotted line: cleavage rate of the matched target; dashed line: limit of detection for the slowest-cleaving targets. Error bars: 5% and 95% confidence intervals as measured by bootstrap analysis. **(c)** ΔABAs (upper, grays) and cleavage rates (lower, blues) for DNAs containing two mismatches. Black boxes expanded in callouts. **(d)** dCas9 ΔABAs (upper) and Cas9 cleavage rates (lower) for DNAs containing a single nucleotide deletion or **(e)** a single nucleotide insertion. Error bars: ΔABA SD and cleavage rate 95% confidence interval as measured by bootstrap analysis. **(f)** Normalized NGS reads for target (TS) and nontarget (NTS) strands of DNAs containing either a mismatch at nucleotide 3 (C3T or A3T) or a deletion at position 1 compared to sgRNA 1 (left) or sgRNA 2 (right). Error bars: maximum SD for cut products produced from cleavage of matched DNA controls. **(g)** Average cut site positions for each strand (TS, NTS) from DNAs containing one mismatch relative to sgRNA 1 (upper) or sgRNA 2 (lower). Range spans the earliest timepoint with >33% cut reads (open diamonds) to the final time point (filled diamonds). Diamond size indicates the average cleavage rates of the associated DNAs. Dashed and solid horizontal lines indicate average cut site positions for matched DNA (M) at early and late time points.

Comparison of wtCas9 binding affinities and cleavage rates for library members harboring single mismatches revealed key wtCas9 characteristics (Figure 2A, 2B and S2A). Targets with an NGG PAM were cleaved rapidly, while DNAs with NGA and NAG PAMs were cleaved much slower, nearing our detection limit (*k_c_* > 10^−5^ sec^−1^)(Jiang et al., 2013; Kleinstiver et al., 2015; Sternberg et al., 2014; Zhang et al., 2014). We also observed a ∼9 nt “seed region”, where wtCas9 is particularly intolerant to mispairing. The seed region for wtCas9 has been variously reported to comprise the first eight to twelve nucleotides immediately upstream of the PAM (Hsu et al., 2013; Jiang et al., 2013; Jinek et al., 2012). Most single mismatches in the seed region reduce DNA binding close to that of unmatched DNA and are cleaved at rates near our detection limit. In contrast, DNAs with single mismatches in positions ∼10-17 retain high apparent binding affinity, but cleavage is almost as slow as seed-mismatched targets. Notably, the cleavage rate is dependent on both the base identity and the position of the DNA mismatch (Figure 2B and S2A). DNAs mismatched within the final protospacer nucleotides (∼18-20) confer almost no sequence specificity to Cas9 binding or cleavage (Figure 2A, 2B and S2A) (Zeng et al., 2018). Taken together, these data establish that our integrated platform quantitatively recapitulates known binding and cleavage specificity features of wtCas9.

Two seed mismatches generally reduced DNA binding to background levels and abolished DNA cleavage (Figures 2C and S2B). However, cleavage rates depended on the mismatch identity. For example, a combination of G2A & A6G mismatches retained a relatively fast cleavage rate (0.0017 s^−1^) compared to other mismatch combinations within the seed (Figure 2C callouts). DNAs with a distal and seed mismatch pair show the broadest range of binding affinities and cleavage rates. Surprisingly, rG-dT mismatches were highly tolerated by wtCas9 (Figure 2C, also see Figure 5). The rG-dT mismatch is a thermodynamically stable wobble interaction that may also form transient Watson-Crick-like mispairs (Kimsey et al., 2015). Within the Cas9 R-loop, rG-dT mismatches are the most stable, while other known non-Watson-Crick interactions (rU-dG and rG-dG) are not well-tolerated, indicating that the protein exerts additional constraints on RNA/DNA duplex formation (Sugimoto et al., 2000).

DNA targets harboring insertions and deletions (indels; relative to the sgRNA/crRNA) have not been comprehensively profiled for any CRISPR nuclease, although a previous study indicates that DNAs containing indels are occasionally cleaved faster than fully-matched DNAs (Lin et al., 2014). We analyzed wtCas9 binding and cleavage of DNAs containing up to two indels compared to the fully-matched DNA (Figure 2D, 2E, S2C and S2D). wtCas9 activity was highly sensitive to deletions in the target DNA. A single deletion in the seed reduces apparent binding affinity and cleavage rates to near-background levels. DNAs with deletions at most PAM-distal positions are bound with intermediate affinity (ΔABAs of 0.2-0.5), while their cleavage rates are ∼100-fold slower than matched DNA (*k_c_* ∼ 10^−3^ s^−1^) (Figures 2D and S2C). Cleavage of DNAs containing two deletions only occurred if positioned within nucleotides 18-20, which rarely affects binding or cleavage (Figure S3A). wtCas9 activity was similarly sensitive to insertions in the target DNA However, DNA targets with insertions at PAM-distal positions 19 & 20 bound DNA with slightly higher affinity than the matched DNA target (Figure S3B). One explanation is that nucleotides upstream of the protospacer weakly alter interactions with Cas9 (as observed by (Kim et al., 2017b)) and programmed insertions reveal their influence. In contrast to DNA binding, cleavage rates were much more sensitive to the identity of the inserted nucleotides (Figures 2E, S2D, and S3B). This difference between DNA binding and cleavage may stem from the R-loop-dependent rearrangement of Cas9 nuclease domains for DNA cleavage (Chen et al., 2017). wtCas9 has lower affinity for DNAs containing two insertions or two deletions and cleaves them slower than DNAs with mismatches in the same positions (Figures 2 and S3), indicating that indels encounter additional steric constraints within the propagating R-loop.

### Analysis of wtCas9 cleavage products

We bioinformatically identified the 5’ ends of target and non-target strands (TS, NTS) of each DNA library member via the unique barcodes on either side of the DNA cut. The expected blunt cut was observed when wtCas9 was charged with sgRNA 1 (Figure 1F, left). However, wtCas9 charged with sgRNA 2 initially produced a one-nucleotide 5’ overhang (Figure 1F, right). After approximately 15 minutes, the 5’ overhang on the NTS recedes, likely due to a second RuvC domain-catalyzed cleavage (trimming). The variability in cleavage products and subsequent trimming activity extended across both DNA libraries. For example, a mismatch in the third position produced a blunt DNA with sgRNA 1, but a trimmed overhang with sgRNA 2 (Figure 2F). Surprisingly, the HNH domain consistently cleaved between the third and fourth bases for all types of single edits, whereas the RuvC domain produced variable overhangs that were strongly dependent on the position and type of mispair (Figure 2G, S2E and S2F). Edits near the scissile bond (between nucleotides 3 and 4) alter the cleavage position and trimming rates (Figure 2G). For example, a mismatch biases wtCas9 towards a blunt cut, presumably through modulation of RuvC domain-mediated NTS cleavage and is even stronger for DNAs harboring a deletion (Figure S2E). Conversely, wtCas9 cleaved DNAs with an insertion at this position further from the PAM (TS +1, NTS +2) (Figure S2F). In sum, the RuvC domain repositions the cleavage site in response to a partially mismatched sgRNA-DNA complex and this behavior is sgRNA and mismatch specific.

### Systematic off-target profiling of engineered Cas9 nucleases

We selected three engineered Cas9 variants (<u>enh</u>anced eSp1.1, Cas9-Enh; high fidelity Cas9-HF1, hyper accurate Cas9-Hypa) for comprehensive binding and cleavage comparison to wtCas9 (Figure 3). As expected, all engineered nucleases retain an NGG PAM. Remarkably, engineered Cas9 variants bound partially-matched DNA libraries with similar affinities to those of dCas9 (r = 0.96, 0.99, 0.96, respectively) (Figures 3A). This result is striking because Cas9-HF1 and Cas9-Enh were both designed to destabilize non-specific Cas9-DNA interactions and were speculated to reduce both off-target DNA binding and cleavage (Kleinstiver et al., 2016a; Slaymaker et al., 2016). In contrast to their DNA binding specificity, all three enzymes showed increased cleavage specificity relative to wtCas9 (Figure 3B). Cas9-Enh, Cas9-HF1, and Cas9-Hypa all rapidly cleave matched DNA (∼0.1 s^−1^); >40% of the partially-matched DNA library was cleaved more slowly by the engineered enzymes than by wtCas9 (Figure 3B). Cas9-HF1 reduced mismatched DNA cleavage rates the most relative to the matched DNA cleavage rate (i.e., increased specificity), followed closely by Cas9-Hypa, then Cas9-Enh, and finally wtCas9.

**Figure 3:**
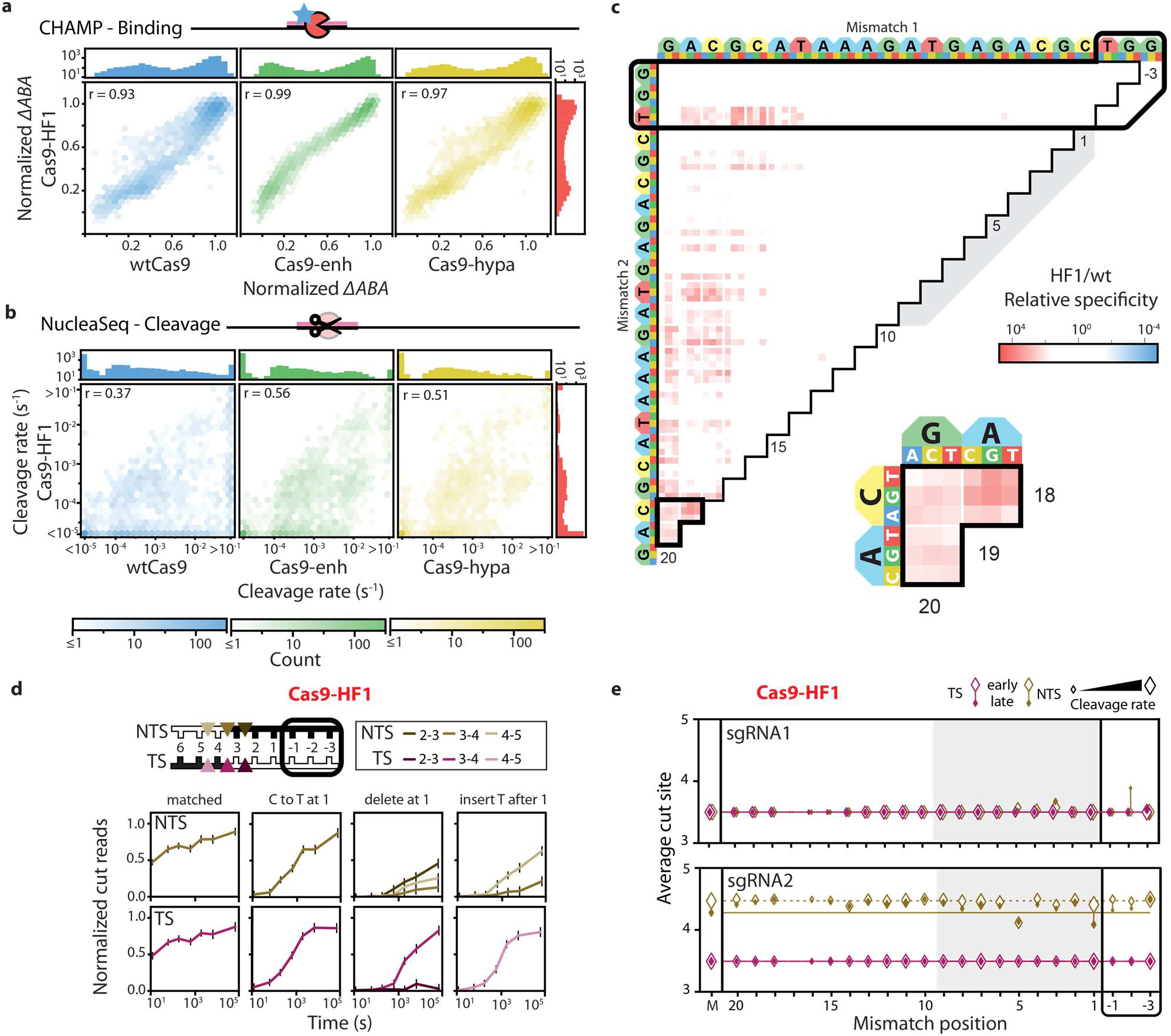
Comparison of engineered Cas9 nucleases. **(a)** Two-dimensional density plots show the correlation of Cas9-HF1 ΔABAs and **(b)** cleavage rates to those from wtCas9, Cas9-enh and Cas9-hypa (sgRNA 1). Top and right histograms include all ΔABAs or cleavage rates for the respective nuclease. Pearson correlation coefficient shown in each panel. Color bars indicate the density of points in each bin. **(c)** Ratio of Cas9-HF1 to wtCas9 specificities for all DNAs containing two mismatches (sgRNA 1). Red: slower cleavage by Cas9-HF1; blue: slower cleavage by wtCas9. Black-outlined range expanded in call-out. **(d)** Cas9-HF1 cleavage patterns on the target (TS) and nontarget (NTS) strands of select DNA targets (sgRNA 1). Normalized counts of cut products comprising ≥15% of the total cut reads are shown. Error bars: maximum SD for any cut product produced from cleavage of the related matched DNA controls. **(e)** Average cut site positions generated by Cas9-HF1 for each strand (TS, NTS) for targets containing a single mismatch relative to sgRNA 1. Range spans the earliest timepoint with >33% cut reads (open diamonds) to the final time point (filled diamonds). Diamond size indicates the average cleavage rates of the associated DNAs. Dashed and solid horizontal lines indicate average cut site positions for matched DNA (M) at early and late time points.

Cas9-HF1 shows the most improved DNA cleavage specificity for targets in the three PAM-distal protospacer nucleotides (positions 18-20) (Figure 3C). Improved specificity was observed for all types of base substitutions and for insertions or deletions at nucleotides 18-20 (Figure S4). Surprisingly, Cas9-HF1 also trims overhangs more slowly than wtCas9; we rarely observed Cas9-HF1 trim DNAs with mismatches, deletions or insertions (Figures 3E and S4C,D). One notable exception occurred when DNAs contained insertions or mismatches near cleavage sites (nucleotides 1-5) (Figures 3D and S4C,D). As with wtCas9, these DNAs still altered both the cleavage site and trimming rate. Different edits elicit different end trimming kinetics. For instance, a C_1_ to T_1_ mismatch (C1T) produces blunt cuts like a matched DNA, but a deletion produces at least three NTS and two TS cleavage products. An insertion at that site shifts the cleavage pattern one nucleotide away from the PAM (Figure 3D) (Liu et al., 2019). In sum, Cas9-HF1 provides the greatest cleavage specificity and the least trimmed DNA ends among the tested Cas9s.

### Profiling *As*Cas12a cleavage specificity

*As*Cas12a is a type V CRISPR-Cas nuclease that is reported to be more specific than wtCas9 in cells (Kim et al., 2016; Yan et al., 2017; Zetsche et al., 2015). We recently showed that the specificity of wtCas12a is due to reversible R-loop propagation, even when the DNA has a PAM-distal mismatch (Strohkendl et al., 2018). Here, we expand our prior work to assay wtCas12a cleavage rates on two DNA libraries, each comprised of >10^5^ partially mispaired DNA targets that provide a direct comparison to engineered Cas9 nucleases (Figure 4, Table S2). As with wtCas9, we recovered the canonical 5’-TTTV PAM (Figure 4A), though wtCas12a cleaved matched DNA 10 to 30-fold more slowly than wt Cas9 (Figure 4B left). Mismatches in the first eight PAM-proximal positions slowed cleavage at least 100-fold, while single PAM-distal mismatches through position 16 were <10-fold slower (Figure 4A). The cleavage rate of mismatched DNAs strongly depended on mismatch identity. For example, a C_17_ to G_17_ mismatch was cleaved twenty-five times more slowly than a C_17_ to T_17_. However, a single mismatch in positions 18-20 did not impact cleavage, as seen with wtCas9. Importantly, cleavage of single-mismatched DNAs correlate strongly with our previously-measured R-loop propagation rates (r=0.91, Figure 4B right) (Strohkendl et al., 2018). Taken together, our work shows that Cas12a cleavage specificity is dominated by rate-limiting R-loop propagation followed by a rapid DNA cleavage step.

**Figure 4:**
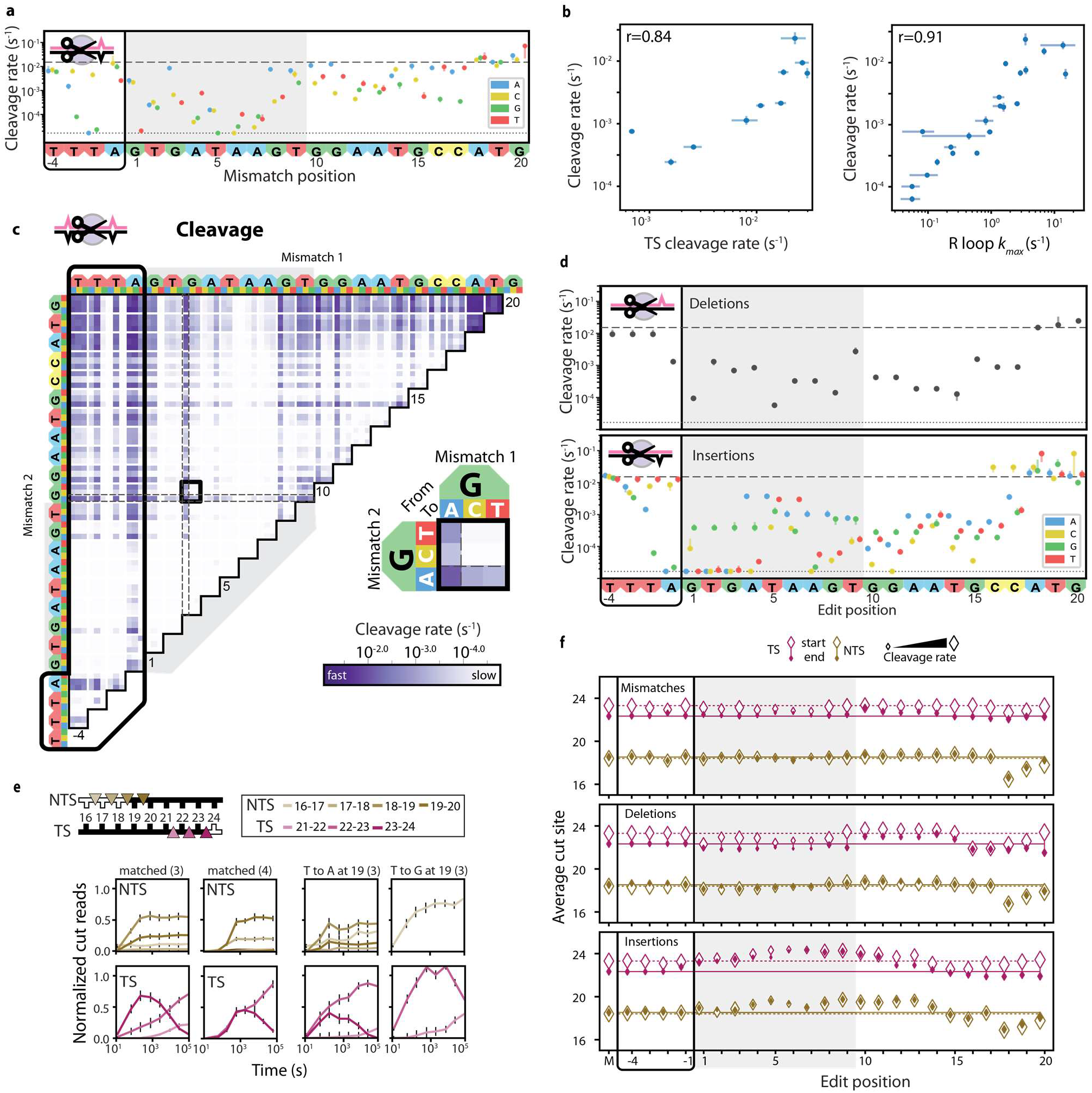
Comprehensive analysis of off-target Cas12a cleavage. **(a)** Cas12a cleavage rates for DNAs containing a single mismatch relative to crRNA 3. Dotted line: cleavage rate of the matched target; dashed line: limit of detection for the slowest-cleaving targets. Error bars: 90% confidence interval as measured by bootstrap analysis. **(b)** Target strand cleavage rate compared to total cleavage rate (left); R-loop propagation (right) is the rate-limiting step for cleavage of mismatched DNA targets. Cleavage rate error bars: 90% confidence interval as measured by bootstrap analysis. TS cleavage rates and R-loop propagation rates with errors bars (SD of three replicates) are taken from (Strohkendl et al., 2018). **(c)** Cleavage rates for DNAs containing two mismatches. Black box expanded in callout. **(d)** Cleavage rates for DNAs containing a single nucleotide deletion (upper) or a single nucleotide insertion (lower). Error bars: Cleavage rate 95% confidence intervals as measured by bootstrap analysis. **(e)** Normalized NGS reads for target (TS) and nontarget (NTS) strands of the indicated DNAs. Parenthesis: corresponding crRNA. Error bars: maximum SD for cut products produced from cleavage of matched DNA controls. **(f)** Average cut site positions for each strand (TS, NTS) from DNAs containing one mismatch (upper), deletion (middle), or insertion (lower) relative to crRNA 3. Range spans the first timepoint with >33% cut reads (early, open diamonds) to the final time point (end, filled diamonds). Diamond size indicates the average cleavage rates of the associated DNAs. Dashed and solid horizontal lines indicate average cut site positions for matched DNA (M) at early and late time points.

We next analyzed wtCas12a cleavage rates of DNAs harboring up to two mismatches (Figure 4C). Two mismatches within the first fourteen PAM-proximal nucleotides typically reduced DNA cleavage rates below our detection limit of <10^−5^ s^−1^ (i.e., for a DNA substrate with G3T & G7C substitutions, Figure 4C callout). In contrast, DNAs with one or two PAM-distal mismatches (at positions 15-20) were cleaved similarly to their related single-mismatched DNAs (vertical banding in Figure 4C). Surprisingly, as with wtCas9, rG-dT mismatches show elevated cleavage rates relative to other types of mismatches at the same positions (Figure 4C, dashed boxes). We did not observe this mismatch preference in the PAM of either nuclease, indicating that both enzymes have similar preferences for stabilizing specific mispairs within a propagating R-loop.

How Cas12a treats DNAs with deletions or insertions relative to the crRNA is unknown, but structural studies suggest that some R-loop bulges may be accommodated within the protein (Gao et al., 2016; Stella et al., 2018; Yamano et al., 2016). A deletion within the first 17 positions slows DNA cleavage >10-fold, while the final positions have no affect (Figure 4D). DNAs with insertions at 1-17 are also slowly cleaved, often beyond the NucleaSeq detection limit. However, cleavage rates vary widely with the identity of the inserted nucleotide (Figure 4D, see 3A4 vs. 3C4), possibly reflecting protein-specific stabilization of the inserted DNA base. Like mismatches and deletions, DNAs with insertions at the final three positions (18-20) do not reduce cleavage rates). These data confirm that pairing with the final crRNA nucleotides is largely dispensable for wtCas12a cleavage, but not overhang trimming (see below).

### Analysis of AsCas12a cleavage products

Cas12a produces staggered 5’ DNA overhangs when it cleaves a crRNA-matched DNA substrate (Zetsche et al., 2015). On matched DNA, we previously showed that the initial TS cleavage site varies by one nucleotide and the NTS overhang is progressively trimmed with time (Strohkendl et al., 2018). Using NucleaSeq, we expand this analysis across two different DNA libraries. wtCas12a produces ∼5-nt 5’ overhangs when cleaving a matched DNA (crRNA 3; Figure 4E). Since NucleaSeq reports on the 5’ strand of cleaved DNAs, we observed that the first NTS cleavage site also varies by one nucleotide. The TS was initially cleaved between nucleotides 23 and 24, but the distribution of cut products changed over time, indicating progressive 5’ trimming, as previously observed for the NTS. Loaded with crRNA 4, wtCas12a still produces ∼5-nt 5’ overhangs, however the NTS cleavage sites vary less and wtCas12a trims the TS overhang more slowly (Figure 4E). After binding the matched target DNA, some Cas12a variants can also nick both single-stranded and double-stranded DNAs *in trans* via a non-specific nuclease activity (Chen et al., 2018; Li et al., 2018; Murugan et al., 2019; Swarts and Jinek, 2018). We monitored Cas12a *trans* cleavage activity by mapping the time-dependent depletion of non-specific DNAs that are not complementary to the crRNA. Second, we looked for the accumulation of truncated DNA fragments that are expected to occur from multiple random Cas12a-generated nicks (Figure S6). Both approaches indicate that *trans* cleavage is minimal under the single turnover conditions employed in this assay.

Cas12a typically produces a similar spectrum of cleavage products on matched and mismatch-containing DNAs (Figure 4F). A notable exception is when the mismatches occur near the NTS cleavage site (nts 18-20) (Figure 4F, top). For example, Cas12a cleaves the NTS up to three nucleotides closer to the PAM when a mismatch is located at the 18^th^ position. The identity of the mismatched nucleotide strongly influences both the cleavage site and the extent of trimming (Figure 4E, right graphs). An rT-dT mismatched DNA (T19A) is cleaved anywhere between nucleotides 16 and 20 in the NTS. However, the NTS of an rT-dC mismatched DNA (T19G) is cleaved between nucleotides 17 and 18 only, reducing the diversity of overhangs even more than matched DNA. TS trimming is also strongly affected; time-dependent changes in cleavage outcomes suggest that wtCas12a trims mismatched DNAs more rapidly (T19G > T19A > matched).

DNAs harboring deletions and insertions deviate from the trends that were observed with mismatched DNA targets (Figure 4F, center). The NTS is processed much like a matched DNA (unless deletions are near the cleavage site). TS cleavage occurs closer to the PAM (i.e., with deletions at nucleotides 4-8 or 16-19) or one nucleotide further away from the PAM (i.e., deletions at nucleotides 9-15). The periodicity of this trend is reminiscent of a full helix turn within the R-loop (10-11 bp) and highlights that a bulge in the crRNA may be more permissible on one face of the helix. For DNAs with insertions, TS and NTS cleavage tracked closely with one another (Figure 4F, bottom). A single insertion at nucleotides 1-14 pushes wtCas12a cleavage of both strands one nucleotide further from the PAM, suggesting that insertions may bulge out of the R-loop to maintain parity with the crRNA. Cleavage sites for DNAs harboring distal indels (18-20) are similar to those with mismatches, suggesting that they are interpreted by wtCas12a as a string of mismatches (compare across Figure 4F Taken together, DNAs containing all three edit types showcase that wtCas12a has considerable flexibility in cleaving both the NTS and TS. This mechanism is consistent with a single RuvC nuclease domain that cuts—and often trims—both strands (Chen et al., 2018; Li et al., 2018).

### Modeling CRISPR-Cas nuclease specificity

To understand the features governing off-target cleavage, we fit the cleavage specificity to several biophysical models of increasing complexity (Figure 5 and Supplemental Methods). In contrast to machine learning-based approaches, these models infer the penalties for off-target cleavage via a limited set of biochemically intuitive parameters. For each nuclease, the models were trained on the entire dataset that includes two distinct DNA target libraries. Training the models on multiple DNA libraries was essential for properly constraining the fit and for obtaining a more comprehensive biochemical description of the enzymes. All models describe PAM recognition via a position weight matrix (PWM) (Stormo and Zhao, 2010). The Cas9 PWM reproduces the canonical NGG and also reflects Cas9’s tolerance of G→A mismatches. Similarly, the model accurately reproduces the Cas12a TTTV PAM with increasing overall specificity from positions −4 to −2 and a modest tolerance for T → C PAM substitutions (Figure S7). Each model is defined by how mismatches and indels within the R-loop are used to calculate the cleavage specificity. The optimal model was selected because it minimizes lost information as defined by the Akaike Information Criterion (AIC), and it accurately captures the variance in the experimental cleavage rates (Akaike, 1974).

**Figure 5:**
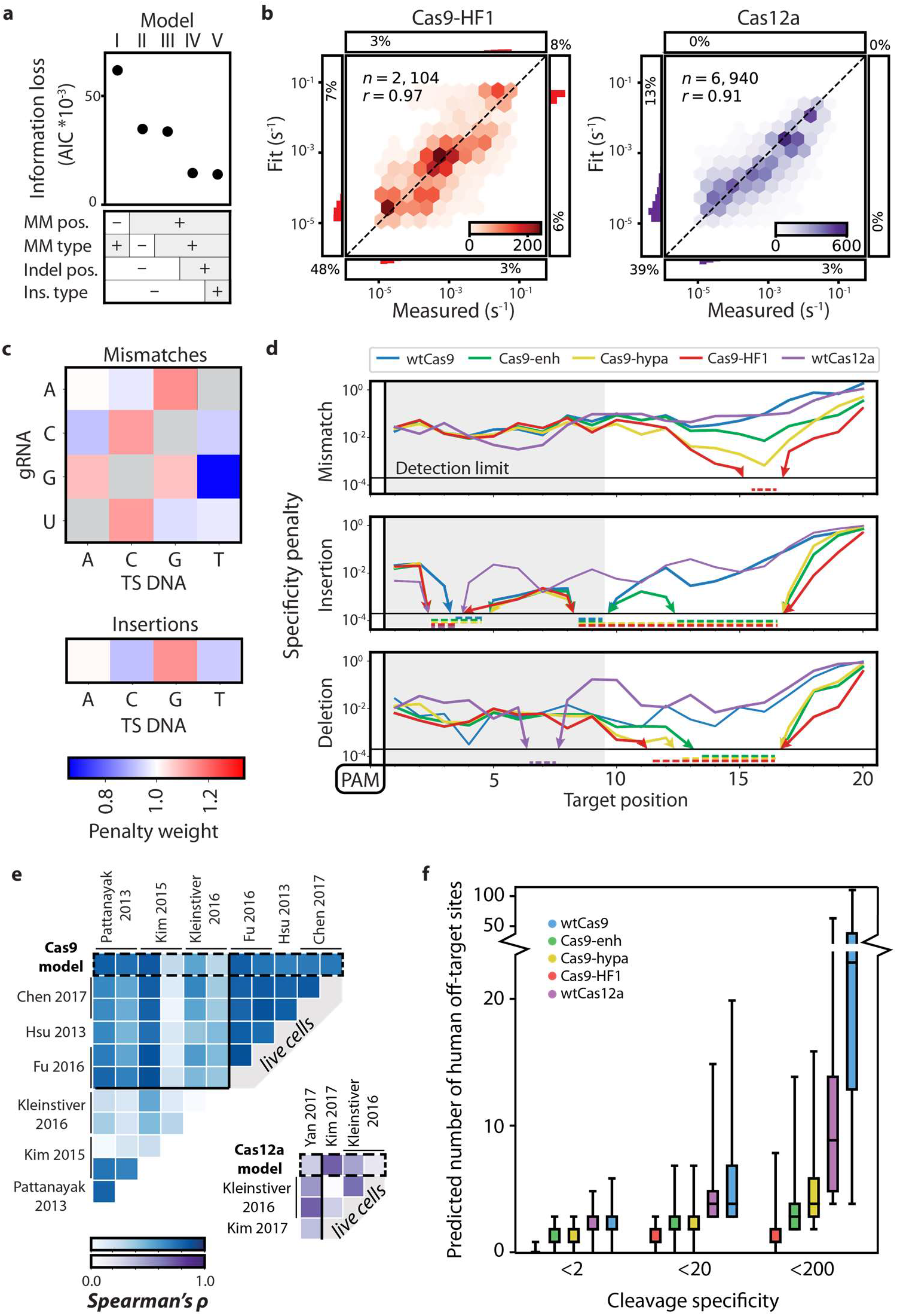
Statistical modeling of CRISPR-Cas nuclease cleavage. **(a)** Akaike Information Criterion (AIC) values for five increasingly detailed biophysical models. The most detailed model has the lowest AIC, i.e. the best goodness of fit. The largest model improvements correspond with addition of R-loop position-specific parameters. **(b)** Correlation between measured and modeled cleavage rates for Cas9-HF1 (left, red) and wtCas12a (right, purple). Color bars indicate the density of points in each bin. Side histograms show the distributions of fit for measured values beyond the upper and lower detection limits. Percentages show the quantity of data in each category with one or both values beyond detection. r = Pearson correlation coefficient. **(c)** Base-dependent weights for mismatches and insertions averaged across all Cas9 and Cas12a enzymes. See Figure S7 and text for additional information. **(d)** Modeled specificity penalties for a single mismatch (upper), insertion (middle), or deletion (lower). All protein variants are oriented with the PAM on the left for comparison. Arrows and dashed lines indicate values below the detection limit (dotted line). **(e)** The predicted reduction in mismatch-dependent cleavage rates is strongly correlated with prior high-throughput measurements of reduced edit efficiencies for wtCas9 (blue) and wtCas12a (purple) (Spearman correlation). See supplementary materials for comparison methodology and associated data. **(f)** The number of off-target sites in the human genome with a predicted cleavage specificity greater than the indicated specificity threshold, shown for each nuclease. For each enzyme, one thousand targets were selected randomly from exomic DNA; the cleavage specificities of the potential off-target cleavage sites across the genome were calculated using model V.

The simplest model assigned a position-independent penalty for each of the twelve types of possible mismatches, regardless of where they occur within the R-loop (Figure 5A, first model). Insertions and deletions were treated as long strings of mismatches. This model only correlated to the measured specificity constants for all five enzymes with correlation coefficients between 0.60 and 0.68 (Figure S7). Including a position-dependent mismatch penalty (but not mismatch type) significantly improved the AIC, highlighting that the position of a mismatch within the R-loop is a critical determinant of the cleavage rate (Figure 5A, second model). The third model also captured the experimental observation that different types of mismatches at a given R-loop position significantly altered Cas9 and Cas12a cleavage rates (e.g., Cas9 tolerates rG-dT mismatches more than rC-dC, Figure 2B). This model combines the position-dependent penalties with an additional position-independent penalty for all twelve possible mismatches. Although the third model improved correlation with measured values with r = 0.86-0.92 (Figure S7), it did not capture how indels altered cleavage specificity. Therefore, the most successful model also included additional position-dependent penalties for insertions and deletions, as well as a position-independent term for each of the four inserted bases (i.e., insertion of a dT vs dA anywhere along the R-loop) (model V, Figure 5B,). These additions further improved the AIC and Pearson’s correlation coefficient, highlighting that each edit has a distinct position-dependent impact on the enzyme’s overall specificity. The final model had the lowest information loss (as shown by the AIC) and had correlation coefficients between 0.93-0.97 with measured cleavage rates for all Cas9 variants and 0.91 for Cas12a (Figure 5B and Figure S7). This model reveals additional biochemical features of each enzyme’s cleavage specificity, as discussed below.

Strikingly, all Cas9 variants and Cas12a have nearly identical mismatched and inserted base preferences (Figure 5C). Mismatch penalties are lowest for rG:dT, rA:dC, rC:dA, rU:dG—basepairs with known wobble and Watson-Crick like mispairing propensity (Sugimoto et al., 1997). However, mismatch penalties derived from thermodynamic studies of RNA-DNA duplexes do not completely recapitulate all Cas9 and Cas12a mispair preferences, indicating that some mismatches are further stabilized within the protein-mediated R-loop (Sugimoto et al., 2000). Insertions showed a reduced penalty for pyrimidines, possibly because they are sterically smaller than purines within a Cas9 or Cas12a-enveloped R-loop. Since both Cas9 and Cas12a tolerated the same types of mismatches and insertions, we conclude that position-dependent specificity penalties are the major determinant of overall differences between the enzymes.

Cas9-HF1 discriminates strongly against all types of PAM-distal edits (Figure 5D, top). For example, Cas9-HF1 cleaves targets with mismatches at positions 14-16 >10^3^-fold slower than the matched target. Mismatches in the 16^th^ position of the R-loop reduce the observed cleavage rates below our detection limit; we therefore report a lower estimate on the actual enzyme specificity (dashed lines in Figure 5D). Cas9-Hypa—originally described as an improved variant of Cas9-HF1—is overall less specific in the PAM-distal region, underscoring the need to comprehensively measure cleavage rates across a large DNA library. Cas9-Enh modestly improves PAM-distal specificity relative to wtCas9 but is still less specific than either Cas9-HF1 or Cas9-Hypa. Insertions and deletions in the PAM-distal region show the largest difference in specificity between WT and the engineered Cas9 variants (Figure 5D, middle, bottom). However, Cas9-HF1 still maintains the highest estimated specificity, with no detectable cleavage of targets with even a single insertion between the 9^th^ and 16^th^ R-loop positions.

In addition to comparing within a class of nucleases, our datasets and biophysical model also allows direct comparison between Cas12a and Cas9. Surprisingly, Cas12a shows a similar cleavage specificity to wtCas9 across the entire R-loop. One minor exception is that Cas12a is more sensitive than wtCas9 to mismatches in positions five through eight. Cas12a also showed lower specificity than wtCas9 for DNAs with indels, with a progressive decrease in specificity as the indel occurs further from the PAM. The penalties for each class of edits point to the same two conclusions: 1) wtCas9 and wtCas12a have similar cleavage specificities, and 2) all engineered Cas9s are more specific than wtCas9, with Cas9-HF1 performing best.

We compared our kinetic model at saturating enzyme concentration to prior *in vitro* and cellular studies of wtCas9 and wtCas12a specificity (Chen et al., 2017; Fu et al., 2016; Hsu et al., 2013; Kim et al., 2015, 2016, 2017a; Kleinstiver et al., 2016b, 2016a; Pattanayak et al., 2013; Yan et al., 2017) (Figure 5E). Importantly, prior cellular high-throughput experiments enumerated off-target sites at multiple loci but at a single time point after transfection. Similarly, prior wtCas9 high-throughput *in vitro* studies also reported a single time point, thereby missing all kinetic and DNA end processing information. To compare different target DNAs, we computed the rank-order (Spearman’s) correlation coefficient between our model and the value for the position-dependent mismatch reported for each of the prior studies’ protospacer sequences. The Spearman correlation determines the strength and direction of the monotonic relationship between two variables that are not linearly related, as occurs between the logarithmically-distributed cleavage rates and the *in vivo* edit efficiencies measured via different reporter assays. (Figure 5E, top). We limited the comparison to studies that measured the effects of mismatches by either changing the DNA or the sgRNA at each position along the R-loop. Two representative genomic targets were selected from studies that included multiple sgRNAs. For wtCas9, our model strongly correlates with both cellular and *in vitro* studies (mean *ρ* = 0.66 ± 0.19; mean ± SD). Interestingly, the correlation between independent datasets is frequently weak (*ρ* = 0.53 ± 0.27), suggesting that the cleavage rates we measured here capture most but not all of the variance in the cellular datasets. The few Cas12a studies that profiled the effect of mismatches along the entire R-loop also had a positive correlation with the model (p = 0.43 ± 0.22; mean ± SD) (Figure 5E, bottom). Using the biophysical model, we next estimated the specificity of each enzyme in the context of the human genome. Figure 5F summarizes the distribution of strong (cleavage specificity < 2), medium (< 20), and weak (<200) off-target genomic sites for one thousand randomly selected exomic targets. Cas9-HF1 has the lowest number of off-target cleavage sites with wtCas9 and wtCas12a showing similar off-target behaviors. The surprising similarity between wtCas9 and wtCas12a *in vitro* suggest that nuclease-extrinsic factors such as the chromatin state, RNP expression level, genomic context, and DNA target sequence may influence wtCas12a cleavage outcomes more strongly than wtCas9 (see Discussion). In sum, NucleaSeq and associated biophysical modeling provides mechanistic insights into enzyme-intrinsic cleavage rates and cleavage products, allows quantitative comparisons between nucleases, and can further improve off-target prediction algorithms.

## Discussion

Newly discovered and engineered nucleases are outstripping our understanding of each enzyme’s cleavage mechanisms and intrinsic specificity. Here, we describe NucleaSeq, which comprehensively profiles cleavage kinetics, cut site structure, and overhang-trimming rates on designer DNA libraries (Figure 1). The DNA library design, post-cleavage analysis, and biophysical modeling software are freely available via GitHub (https://github.com/finkelsteinlab/nucleaseq). Future modifications to NucleaSeq can extend this platform for measuring the kinetics of off-target base-editing, DNA nicking, and CRISPR-transposon DNA insertions (Klompe et al., 2019; Strecker et al., 2019). Similarly, CHAMP can be extended via *in vitro* transcription and translation to measure protein-RNA and protein-peptide interactions (Buenrostro et al., 2014; Ozer et al., 2015).

Using the large NucleaSeq datasets, we develop a biophysical model that facilitates comparison across different types of nucleases, captures intrinsic nucleotide mispair preferences, and reveals additional biochemical features of each enzyme. Modeling cleavage specificity across multiple targets is especially important as we and others have observed that Cas9 and Cas12a have poorly-understood but pervasive target-specific behaviors (Doench et al., 2014; Moreno-Mateos et al., 2015; Wang et al., 2014; Xu et al., 2017). In addition to specifying the appropriate enzyme for a specific gene editing application, the integration of these datasets into off-target prediction servers will improve genomic target DNA selection.

All engineered Cas9 variants profiled here, and most notably Cas9-HF1, retain similar off-target DNA binding affinities but improve cleavage specificity of DNAs with PAM-distal mispairs (Figure 5). However, all Cas9 variants had similar cleavage specificity in the PAM-proximal region. Taken together, our data indicate that this increased specificity is largely driven by slowing the cleavage step after R-loop propagation (e.g., conformational changes that activate the nuclease domain(s)) (Liu et al., 2019). This observation is consistent with the observation that mutations in the *Sp*Cas9 bridge helix can regulate the HNH nuclease domain prior to cleavage (Babu et al., 2019; Nishimasu et al., 2014; Sternberg et al., 2015). Our data also indicates that off-target DNA binding remains an ongoing challenge for Cas9 engineering and for dCas9-based applications (i.e., CRISPRi, CRISPRa, and base-editing) (Gilbert et al., 2014; Komor et al., 2016; Qi et al., 2013). In contrast to *Sp*Cas9, Cas12a has a late transition during R-loop formation, making dCas12a a strong candidate for applications that require high DNA binding specificity (Strohkendl et al., 2018).

Deep characterization of Cas9 and Cas12a reveals shared features that highlight convergence of these phage defense systems. They share similar cleavage specificities and tolerate similar types of mispairs (i.e., rG-dT mismatches and pyrimidine insertions), exchanging fidelity and off-target cleavage in order to target rapidly evolving phages. The RuvC domain of both enzymes can also create staggered cuts and trim DNA overhangs, possibly limiting error-free re-ligation of the invading nucleic acid. Kinetic discrimination against partially matched DNA targets may result from slowed R-loop propagation and/or slowed cleavage after the R-loop is formed. As R-loop propagation is the rate-limiting step for cleavage by both Cas9 and Cas12a, it is the major determinant for enzyme specificity at sub-saturating concentrations that may dominate the cellular environment (Boyle et al., 2017; Gong et al., 2018; Liu et al., 2019; Strohkendl et al., 2018).

*As*Cas12a and related Type V nucleases are reportedly more specific than Cas9 in human cells (Kim et al., 2016; Yan et al., 2017). However, wtCas12a has a remarkably similar *in vitro* cleavage specificity to that of wtCas9 at saturating enzyme concentrations (Figure 5). One possible explanation for the discrepancy between the cellular and *in vitro* observations is that wtCas12a cleaves both matched and off-target DNAs 10-to 40-fold slower than wtCas9. This slower overall cleavage rate may provide sufficient time for wtCas12a to be displaced from off-target genomic sites in cells (i.e., by the transcription apparatus) without changing the *in vitro* specificity. In support of this idea, NmeCas9 was recently reported as high-fidelity and has a much slower cleavage rate than *Sp*Cas9 *in vitro* (Amrani et al., 2018; Edraki et al., 2018). The Cas12a-generated DNA overhang structure can bias the cellular repair pathway towards homologous recombination or non-homologous end joining. We propose that programming specific PAM-distal mismatches between the crRNA and target DNA can direct the repair outcomes without changing the overall cleavage rate (i.e., genome editing efficiency) for both Cas12a and Cas9.

In sum, the integrated NucleaSeq and CHAMP platform provides a framework for profiling and engineering high-fidelity nucleases for gene editing applications. Engineered Cas9 variants generally improve specificity at a narrow set of R-loop positions but the relatively low seed-region mismatch specificity and inconsistent cut site structure across different target sequences are areas for further improvement. Future enzyme engineering efforts should aim to combine Cas12a-like binding specificity with Cas9-like conformational cleavage regulation. We anticipate that the integrated NucleaSeq and CHAMP workflow will be broadly useful for guiding enzyme profiling and engineering strategies.

## Materials and Methods

### Oligonucleotides, CRISPR RNA, and DNA libraries

Oligonucleotides were purchased from IDT (see **Table S1**). Single guide RNAs (sgRNAs) for Cas9 and CRISPR RNAs (crRNAs) for Cas12a were purchased from Synthego (see **Table S1**). Pooled oligonucleotide libraries were purchased from CustomArray Inc. and Twist Biosciences (see **Table S2**). Libraries were amplified via 12 cycles of PCR with Phusion polymerase (NEB).

#### DNA Library Design

Each library contains DNAs that are variations of a matched DNA sequence (defined by nuclease PAM preference and RNA guide), termed a ‘modified target’. Modified targets include: single and double substitutions, insertions, or deletions, and all sequences with a contiguous subsection changed to the complementary bases. Each modified target is flanked by the following additional sequence elements necessary for NucleaSeq analysis and (5’ to 3’): left primer, left barcode, left buffer, modified target, right buffer, variable length buffer, right barcode, right primer (Supplemental File 1). As controls, we included 146 copies of the matched target. Each copy had a unique left and right barcode set. Finally, we included 150 pseudo-random barcoded DNA strands to normalize read depth between time points and biological replicates (see below).

Our libraries use unique barcodes appended to either end of each DNA strand (Hawkins et al., 2018). By searching for the barcodes after NGS, any cleaved DNA can be computationally identified from a partial fragment after cleavage. These barcodes are 17 bp, uniquely paired, and are correctly identified despite any combination of up to two substitutions, insertions, or deletions in their sequence. Similarly, primer sequences (common across the library) were selected that help distinguish left barcodes, right barcodes and cleaved ends. They are distinguishable from one another and the cleaved end of any library member cut within 5 bp of a canonical cut site.

Flanking each modified target are left and right 5 bp buffer regions held constant for all sequences to provide a constant local DNA context for nuclease activity. These buffer sequences were randomly generated with nearly equal nucleotide content. Oligos with insertions and deletions also included a variable-length buffer to ensure that these oligos were the same length as the matched target.

### Protein cloning and purification

*S. pyogenes* Cas9 variants were generated via Q5 site-directed mutagenesis (New England Biolabs) of a pET-based plasmid (pMJ806) (Jinek et al., 2012). Nuclease dead Cas9 variants contained the D10A and H840A mutations. Enhanced-, HF1-, and Hypa-Cas9 variants harbored the mutations indicated in **Table S1** (Chen et al., 2017; Kleinstiver et al., 2016a; Slaymaker et al., 2016). An N-terminal 3xFlag epitope was introduced for fluorescent imaging of nuclease dead variants via CHAMP (see below).

Cas9 protein variants were expressed in BL21 star (DE3) cells (Thermo Fisher Scientific) using a previously established protocol with minor modifications (Jinek et al., 2012). A 4L flask containing 1L LB + Kanamycin was inoculated with a single colony and then grown to an optical density (OD) of 0.6 at 30°C with shaking. Protein expression was induced with 1mM IPTG during 18 hours at 18°C with shaking. Cells were collected by centrifugation and lysed by sonication at 4°C in lysis buffer (20 mM Tris-Cl pH 8.0, 250 mM NaCl, 5mM imidazole, 5 μM phenylmethylsulphonyl fluoride, 6 units ml^−1^ DNAse I). The lysate was clarified by ultracentrifugation at 35k RCF, then passed over a nickel affinity column (HisTrap FF 5mL, GE Healthcare) and eluted with elution buffer (20 mM Tris-Cl pH 8.0, 250 mM NaCl, 250 mM imidazole). The His_6_-MBP was proteolyzed overnight in dialysis buffer (20 mM HEPES-KOH pH 7.5, 150 mM KCl, 10% glycerol, 1mM DTT, 1mM EDTA) supplemented with TEV protease (0.5 mg per 50 mg protein). The dialyzed protein was resolved on a HiTrap SP FF 5mL column (GE Healthcare) with a linear gradient between buffer A (20 mM HEPES-KOH pH 7.5, 100 mM KCl) and buffer B (20 mM HEPES-KOH pH 7.5, 1 M KCl). Protein-containing fractions were concentrated via dialysis (10 kDa Slide-A-Lyzer, Thermo Fisher Scientific), and then sized on a Superdex 200 Increase 10/300 column (GE Healthcare) pre-equilibrated into storage buffer (20 mM HEPES-KOH pH 7.5, 500 mM KCl). The protein was snap frozen in liquid nitrogen and stored in 10μL aliquots at −80°C.

*Acidaminococcus sp* (*As*) Cas12a was expressed as an N-terminal His_6_-TwinStrep-SUMO fusion in a pET19-based plasmid (pIF502) (Strohkendl et al., 2018). The Cas12a fusion protein was expressed in BL21 star (DE3) cells (Thermo Fisher Scientific) using a previously established protocol with minor modifications (Strohkendl et al., 2018). A 20 mL culture of Terrific Broth (TB) + 50 mg mL^−1^ carbenicillin was inoculated with a single colony and grown overnight at 37°C with shaking. A 4 L flask containing 1 L of TB was inoculated with 10 mL of the starter culture and then grown to an optical density (OD) of 0.6 at 37°C. Protein expression was induced with 0.5 mM IPTG during 24 hours at 18°C. Cells were collected by centrifugation and lysed by sonication at 4°C in lysis buffer (20 mM Na-HEPES pH 8.0, 1 M NaCl, 1 mM EDTA, 5% glycerol, 0.1% Tween-20, 1 mM PMSF, 2000 U DNase (GoldBio), 1X HALT protease inhibitor (Thermo Fisher)). The lysate was clarified by ultracentrifugation at 35k RCF, applied to a hand-packed StrepTactin Superflow gravity column (IBA Life Sciences), and then eluted (20 mM Na-HEPES, 1 M NaCl, 5 mM desthiobiotin, 5 mM MgCl_2_ and 5% glycerol). The eluate was concentrated to <1mL using a 30 kDa MWCO spin concentrator (Millipore), SUMO protease was added at 3μM, and then the eluate was incubated overnight on a rotator at 4°C.The protein was resolved on a HiLoad 16/600 Superdex 200 Column (GE Healthcare) pre-equilibrated with storage buffer (20 mM HEPES-KOH, 150 mM KCl, 5 mM MgCl2, 2 mM DTT buffer). The protein was finally snap frozen in liquid nitrogen and stored in 10μL aliquots at −80°C.

Cas9 and Cas12a ribonucleoprotein (RNP) complexes were reconstituted by incubating a 2:3 molar ratio of apo protein and RNA (sgRNA and pre-crRNA for Cas9 and Cas12a, respectively) in RNP buffer (20 mM HEPES pH 7.5, 150 mM KCl, 10 mM MgCl_2_, 2 mM DTT) at room temperature for 30 minutes prior to each experiment. Reconstituted RNPs were diluted in the experimental reaction buffer, used immediately, and discarded after the experiment.

### NucleaSeq

DNA libraries were mixed in buffer (20 mM HEPES pH 7.5, 150 mM KCl, 10 mM MgCl_2_, 2 mM DTT) at room temperature with RNP complex to final concentrations of 10 nM and 100 nM, respectively. Aliquots were transferred to a stop solution (final concentration: 12 mM EDTA and 12 U proteinase K (Thermo Fisher)) at the following time points: 0, 0.2, 0.5, 1, 3, 10, 30, 100, 300 and 1000 minutes. The stopped reactions were incubated at 37°C for 30 minutes to remove Cas9 and Cas12a from their DNA substrates. Each time point was ethanol precipitated and resuspended in TE buffer. Samples were submitted to the University of Texas Genomic Sequencing and Analysis Facility, where sequencing adapters (NEBNext Ultra, NEB) were appended. The samples were sequenced on a MiSeq or NextSeq 500 sequencer (Illumina).

#### Bioinformatic analysis pipeline

From each paired-end read pair, we inferred the maximum likelihood full-length sequence using the overlapping base pairs as described previously (Jung et al., 2017). Primer and barcode sequences were then used to identify the intended sequence identity and, for cleaved products, the observed side. Observed and intended sequences were aligned using either global alignment (Cock et al., 2009) for uncleaved products or global alignment with cost-free ends (Cieślik et al., 2016) for cleaved products. Throughout this process, sequences were filtered for quality based on length, primer and barcode structure, and number of synthesis and sequencing errors. Sequences with errors were not allowed in the target and buffer regions.

Next, the read counts for each full-length library member in each sample were normalized to account for two sources of variation. First, we normalized the different total numbers of reads across different time points for each sample. Specifically, each member’s read count for each sample was normalized by the ratio of total read counts at that time point to the total read count of an input control sample (not treated with nuclease). Second, read counts were normalized to account for changes due to sampling from a library of changing composition. The generation of cleaved products and corresponding depletion of full-length products by nuclease activity changes the number of sampled sequences of all species, including species unaffected by the nuclease. To account for this, we used the 150 non-target control sequences as a reference. For each randomly-generated non-target sequence, there is a small probability it will be susceptible to nuclease cleavage. Hence, we used the median read count value of all the random sequences as a robust measure of changes due only to sampling from a library of changing composition (non-target median). Read counts of each library member at each time point were normalized by the ratio of the nontarget median at that time point to the non-target median from the control sample.

In addition to the above two steps, cleaved products were normalized to account for differences in PCR amplification between cleaved products and full-length oligos. We observed that the normalized number of cleaved products should be proportional to the depletion of the corresponding full-length oligos. Stated as an equation, let |*F*|_*t*_ be the number of full-length product reads and |*C*|_*t*_^*side*^ be the number of cleaved product reads on a given side at a given time, for a single library member of choice, normalized as above. Then for normalization and proportionality constants *Z_t_^side^* and *k^side^*,

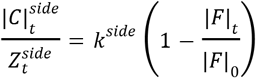

We choose to set the final normalization constant 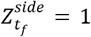 and solve the above for *k^side^*. Plugging this back in and rearranging gives normalization constants

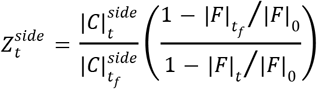

This is intentionally a function only of ratios of read counts, not absolute read counts. This lets us use the median read count ratios from all 146 matched target controls to calculate the normalization constants. These final normalization constants are then used for all library members. Finally, read counts are normalized to range between zero and one. For full-length products, we normalize by the fit value of reads at time zero. For cleaved products, we normalize first by the sum of all cleaved products at all time points, then normalize to set the resulting median sum of all cleaved products at the final time point to the depletion of full-length products, 1 − |*F*|*_t_f__*/|*F*|_0_.

The normalized read counts were fit to a single exponential decay. We observed that the data was well described by a single exponential, implying a constant reaction rate under the single turnover conditions used in this assay. A small fraction of the starting DNA sequences of each species were never cleaved, possibly indicating some hydrolytically inactive enzymes. We thus fit for exponential decay with a constant offset. For the constant offset, we used the median normalized fraction of uncleaved sequences of the 146 perfect target sequences at the final time point. Error bars give the standard deviation of 50 bootstrap measurements, each of which was calculated by resampling the raw read counts with replacement, renormalizing, and refitting (Efron and Tibshirani, 1993).

### Modeling cleavage specificity

#### Model description

We modeled cleavage specificity (*Model 1*), given as the ratio of the cleavage rate of a given sequence *s, k_s_*, to the cleavage rate of the matched sequence *m, k_m_*, as:

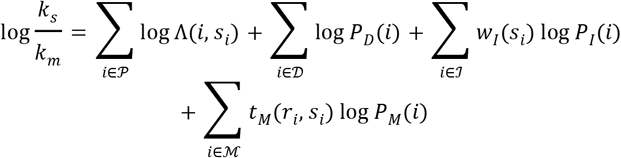

The terms of the model give cleavage rate penalties for the following sequence edits respectively: suboptimal bases in the PAM, target deletions, target insertions, and target mismatches, each with corresponding set of positions with the given sequence edit type: 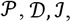, and 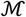. For suboptimal PAM bases, the cleavage rate penalty is given by the function Λ, a function of both the suboptimal base identity, *s_i_*, and its position *i*.

For deletions, insertions, and mismatches, the cleavage rate penalty functions *P_D_, P_I_*, and *P_M_* are dependent only on the position *i*, reflecting the fact that position in the target is the primary determinant of the effect of a given sequence edit. This is intuitive for deletions, as they primarily require steric adjustments to realign the matching base pairs. For mismatches, position was determined to be the primary determinant of the cleavage rate penalty via comparison with other models (see Simplified models below). Insertions have a weighting function *w_I_* to allow for different inserted bases to have different penalties. The base identities in the mismatch are modeled via the weighting function *t_M_*(*r_i_, s_i_*), a function of the mismatched gRNA base *r_i_* and target strand base *s_i_*.

Within the terms for insertion and mismatch penalties, there is an unconstrained degree of freedom in the relative magnitudes of the weights relative to the log position penalties. To remove this extra degree of freedom, the insertion and mismatch weighting functions *w_I_* and *t_M_* were each constrained to have an average value of 1. This was accomplished through the use of Hadamard matrices, possible because *w_I_* and *t_M_* have 4 and 12 parameters, respectively. Hadamard matrices are maximal-determinant matrices using elements of only 1 and −1. We used Hadamard matrices with negative one in all elements not in the first row or column along diagonals 0, −1, 2, −3, −4, −5, 6, 7, 8, −9, and 10, where 0 is the main diagonal and diagonal indices increase up and to the right. We parameterize a constrained length *n* weight vector *w* with a length (*n* − 1) vector *x* of free parameters as follows. Let *H_n_* be the *n×n* Hadamard matrix described above. Due to the inverse identity of Hadamard matrices and the first row and column of *H_n_* being composed entirely of ones, parameterizing with *x* and using the following conversions enforces an average value of 1 in the weights vector *w*:

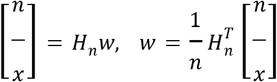

Cleavage rates that are shorter than the first time point or longer than the last one cannot be modeled accurately. We therefore constrained the output of our models with the following “bandpass filter” function:

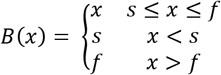

Where *s* and *f* are the slowest and fastest detectible cleavage rates, corresponding to half-lives at our first and last time points.

Ridge regularization of the difference of insertion and mismatch weights from one was used to reduce over-fitting of the underlying cleavage data (Hoerl and Kennard, 1970). Figure S7D shows the fit weight values as a function of the regularization parameter *λ*. The relative parameter values appear to stabilize near *λ* = 10^3^, which we used to fit the model.

#### Simplified models

For comparison, we fit our data to four simplified models, each excluding some terms and/or factors in the full model above. The first three simplified models did not include the insertion or deletion terms, modeling the possibility that the recognition channel does not accommodate bulges to realign matching sequences after indels. Under this assumption, for example, a sequence with a single insertion between the first and second bases, but otherwise perfectly matching, would result in about 75% mismatches due to a forced frame shift. These three models were: cleavage rate as a function of only the mismatch base pair identities, only the mismatch position, or both as in the full model above. The fourth simplified model included insertions and deletions but omitted the insertion weights *w_I_*. Each simplified model included the PAM term. We number the models for reference:

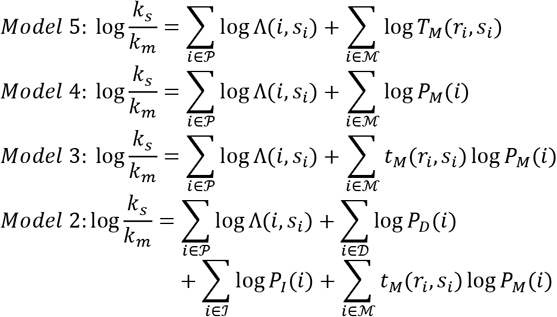

And Model 1 is the full model above. The mismatching base pairs function in Model 5, *T_M_*(*r_i_, s_i_*), is different from the analogous weighting function *t_M_*(*r_i_, s_i_*) in the other models as it gives absolute penalty values, not weights constrained to average value of one.

Figure 5 compares these models using the Akaike Information Criterion (AIC) (Akaike, 1974). The significant improvement in AIC between Models 5 and 4 demonstrates that position is in fact the primary determinant of mismatch cleavage rates. Model 3 demonstrates that including the mismatched base pair identities is a useful but relatively small improvement to the position-only model. Similarly, Models 2 and 1 show that adding insertions and deletions to the model provides a significant improvement, while the addition of insertion weights is a relatively small improvement to the model (i.e., insertions are weakly sensitive to the inserted base identity).

#### Model comparison to previously published datasets

To compare the model’s output with prior measures of nuclease specificity, we selected *in vitro* and *in vivo* published datasets for either *Sp*Cas9 or *As*Cas12a that contained at least one measure of specificity per position in the sgRNA (for *Sp*Cas9) or the crRNA (for *As*Cas12a). Dataset 1 (Pattanayak et al., 2013) included representative genes CLTA1 and CLTA2 with sgRNA v2.1 and 100 nM wt Cas9. Published specificity scores were averaged across all single mismatch values at each position. Dataset 2 (Kim et al., 2015) used Digenome-seq and included sgRNAs targeting genes HBB and VEGFA. Dataset 3 (Kleinstiver et al., 2016a) used GUIDE-Seq to profile indels at genes VEGFA-2 and EMX1-1. Values were extracted from the published heatmaps based on RGB values as measured with FIJI (Schindelin et al., 2012). The measured scores were averaged across all single mismatch values at each position. Dataset 4 (Fu et al., 2016) *in vivo* log retention scores for genes UNC-22A and ROL6 were extracted from published graphs with a data digitization tool (Rohatgi, 2019). The measured scores were averaged across all single mismatch values (transitions and transversions) at each position. Dataset 5 (Hsu et al., 2013) used SURVEYOR nuclease to determine mean cleavage results for aggregated EMX1 targets. Values were extracted from the published heatmaps based on position-averaged RGB values as measured with FIJI (Schindelin et al., 2012).

Dataset 6 (Chen et al., 2017) used a T7E1 reporter assay and included representative genes FANCF-1 and FACNF-4. Percent of modification for each gene was extracted from the published heatmaps based on RGB values as measured with FIJI for wtCas9 (Schindelin et al., 2012). Dataset 7 (Yan et al., 2017) used BLISS to generate composite mismatch tolerances for each guide position. Values were extracted from the published graph with a data digitization tool (Rohatgi, 2019). Dataset 8 (Kim et al., 2017) relative indel frequency values at each position were extracted from the published graph with a data digitization tool (Rohatgi, 2019). Dataset 9 (Kleinstiver et al., 2016b) used a T7E1 reporter assay and included representative gene DNMT1, sites 1 and 3. Percent of modification for each gene was extracted from the published graphs with a data digitization tool (Rohatgi, 2019). Since the measure and distribution of data varied from study to study, a nonparametric correlation was used (only requires ordinal data). Each dataset was compared to one another and to our model’s average positional mismatch penalty to generate Spearman’s rank correlation coefficients (*ρ*). The average mismatch penalty is denoted as *P_M_* in *Model 1*.

### CHAMP (Chip Hybridized Association Mapping Platform)

DNA libraries were sequenced on a MiSeq using 2×75 paired-end chemistry (v3, Illumina) Sequenced MiSeq chips were stored at 4°C in storage buffer (10 mM Tris-Cl, pH 8.0, 1 mM EDTA, 500 mM NaCl) until needed for CHAMP.

Chips were regenerated similarly to our previous strategy (Jung et al., 2017). Each chip was loaded into a custom microscope stage adapter, with temperature controlled by a custom heating element. All solutions were pumped through the chip at 100 μl min^−1^ using a syringe pump (Legato 210, KD Scientific), with reagents added via an electronic injection manifold (Rheodyne MXP9900). Chip DNAs were made single-stranded with 500 μl 60% DMSO, then washed with 500 μl TE buffer. An unlabeled regeneration primer (user DNA specific) and a digoxygenin labeled primer (PhiX DNA specific, for alignment) were annealed over an 85-40°C temperature gradient (30 min) in hybridization buffer (75 mM tri-sodium citrate, pH 7.0, 750 mM NaCl, 0.1% Tween-20), and then excess primers were removed at 40°C with 1 ml wash buffer (4.5 mM Trisodium Citrate, pH 7.0, 45 mM NaCl, 0.1% Tween-20). Annealed primers were extended at 60°C using 0.08 U μl^−1^ Bst 2.0 WarmStart DNA polymerase (New England Biolabs) and 0.8 mM dNTPs in isothermal amplification buffer (20 mM Tris-HCl, pH 8.8, 10 mM (NH_4_)_2_SO_4_, 50 mM KCl, 2 mM MgSO_4_, 0.1% Tween-20), then washed with 500 μl wash buffer. Using 100 μl of rabbit anti-digoxigenin monoclonal antibody (Life Technologies) and 100 μl Alexa488-conjugated, goat anti-rabbit antibody (Thermo Fisher Scientific), PhiX DNA clusters were fluorescently labeled as markers for subsequent image alignment. The MiSeq chips were imaged on a Ti-E microscope (Nikon) in a prism-TIRF configuration (Jung et al., 2017). Images were acquired in OME-TIFF format (uncompressed TIFF plus XML meta data) using the Micro-Manager software (Edelstein et al., 2014).

The dCas9/sgRNA RNP complex was diluted to concentrations of 0.1, 0.3, 1, 3, 10, 30, 100 and 300 nM in CHAMP buffer (20mM Tris-HCl pH 7.5, 100 mM KCl, 5 mM MgCl_2_, 5% glycerol, 0.2 mg mL^−1^ BSA, 0.1% Tween-20, 1 mM DTT). Starting with the lowest concentration, 100 μl of RNP complex was injected into the regenerated MiSeq chip at room temperature and incubated for 10 minutes. Then, 300 μl of CHAMP buffer containing 4 nM Alexa488-conjugated anti-Flag antibody (Alexa Fluor 488 antibody labeling kit, Thermo Fisher; monoclonal BioM2, Sigma-Aldrich) was injected to wash off unbound RNP and label DNA-bound RNP complex. The chip was then imaged over 420 fields of view (FOVs) with 10 frames of 50 ms each, while illuminated with 10 mW of laser power, as measured at the front face of the prism. Collected images were processed via the CHAMP bioinformatic software for downstream analysis (Jung et al., 2017).

### DNA Radiolabeling and PAGE purification

DNA oligonucleotides 308 and 310 were 5’-radiolabeled with [γ-^32^P] ATP (Perkin-Elmer) using T4-polynucleotide kinase (New England Biolabs). Radiolabeled nucleotides were then purified by electrophoresis in a 12% native polyacrylamide gel, before being eluted in TE buffer (10mM Tris-HCl pH 8, 1 mM EDTA) to a concentration 250nM.

### Nuclease active site titration

Atto647N-labeled target DNA was generated with Q5 DNA polymerase (NEB) using oligonucleotides 365, 460 and 371. The DNA was diluted in series from 512 nM to 4nM in reaction buffer (20 mM HEPES pH 7.5, 150 mM KCl, 10 mM MgCl_2_, 2 mM DTT). RNP complexes were formed by mixing protein and RNA (256nM:384nM) and incubating for 30 minutes at room temperature in the same buffer conditions. Equal volumes of RNP and Atto647N-labeled matched DNA dilutions were combined then incubated for 30 minutes at room temperature. The reaction was stopped by addition of a stop solution (40 mM EDTA and 50 U proteinase K (Thermo Fisher)) and a 30-minute incubation at 37°C removed RNPs from their DNA substrates. All samples were run in a 10% polyacrylamide native gel and then imaged using a Typhoon FLA9500 gel scanner (GE Healthcare).

## Author Contributions and Notes

SKJ, JAH, CJ, JRR, WHP, and IJF designed research. SKJ, NVH, KH, CJ and JSC performed research. JAH, JRR, KH, WHP wrote the software. JAH, SKJ and KH analyzed the data. SKJ, JAH, and IJF wrote the paper with editorial assistance from all authors.

This article contains supporting information online.

## Conflicts of Interest

JD is a co-founder of Caribou Biosciences, Inc., Editas Medicine, Intellia Therapeutics, Scribe Therapeutics, and Mammoth Biosciences. JD is a scientific advisory board member of Caribou Biosciences, Inc., Intellia Therapeutics, eFFECTOR Therapeutics, Scribe Therapeutics, Synthego, Mammoth Biosciences, and Inari. JD is a member of the board of directors at Driver and Johnson & Johnson and has sponsored research projects by Roche Biopharma and Biogen. JC is a co-founder of Mammoth Biosciences. The remaining authors declare that the research was conducted in the absence of any commercial or financial relationships that could be construed as a potential conflict of interest.

## Acknowledgments

We thank Isabel Strohkendl, Rick Russell, and members of the UT-Austin Genomic Sequencing and Analysis Facility staff for valuable insights. We are grateful to members of the Finkelstein laboratory for carefully reading the manuscript, and for additional contributions by Kaylee Dillard, Fatema Saifuddin and Jacob Kula. This work was supported by a College of Natural Sciences Catalyst award, the Welch Foundation (F-1808 to I.J.F.), and the National Institute of Health (R01GM124141 to I.J.F.; F32 AG053051 to S.K.J.).

## Notes

https://github.com/finkelsteinlab/nucleaseq

## References

Akaike, H. (1974). A new look at the statistical model identification. IEEE Trans. Autom. Control 19, 716–723.

Amrani, N., Gao, X.D., Liu, P., Edraki, A., Mir, A., Ibraheim, R., Gupta, A., Sasaki, K.E., Wu, T., Donohoue, P.D., et al. (2018). NmeCas9 is an intrinsically high-fidelity genome-editing platform. Genome Biol. 19, 214.

Anderson, K.R., Haeussler, M., Watanabe, C., Janakiraman, V., Lund, J., Modrusan, Z., Stinson, J., Bei, Q., Buechler, A., Yu, C., et al. (2018). CRISPR off-target analysis in genetically engineered rats and mice. Nat. Methods 15, 512.

Babu, K., Amrani, N., Jiang, W., Yogesha, S.D., Nguyen, R., Qin, P.Z., and Rajan, R. (2019). Bridge Helix of Cas9 Modulates Target DNA Cleavage and Mismatch Tolerance. Biochemistry 58, 1905–1917.

Boyle, E.A., Andreasson, J.O.L., Chircus, L.M., Sternberg, S.H., Wu, M.J., Guegler, C.K., Doudna, J.A., and Greenleaf, W.J. (2017). High-throughput biochemical profiling reveals sequence determinants of dCas9 off-target binding and unbinding. Proc. Natl. Acad. Sci. 114, 5461–5466.

Buenrostro, J.D., Araya, C.L., Chircus, L.M., Layton, C.J., Chang, H. Y., Snyder, M.P., and Greenleaf, W.J. (2014). Quantitative analysis of RNA-protein interactions on a massively parallel array reveals biophysical and evolutionary landscapes. Nat. Biotechnol. 32, 562–568.

Chen, J.S., Dagdas, Y.S., Kleinstiver, B.P., Welch, M.M., Sousa, A.A., Harrington, L.B., Sternberg, S.H., Joung, J.K., Yildiz, A., and Doudna, J.A. (2017). Enhanced proofreading governs CRISPR–Cas9 targeting accuracy. Nature 550, 407–410.

Chen, J.S., Ma, E., Harrington, L.B., Costa, M.D., Tian, X., Palefsky, J.M., and Doudna, J.A. (2018). CRISPR-Cas12a target binding unleashes indiscriminate single-stranded DNase activity. Science 360, 436–439.

Crosetto, N., Mitra, A., Silva, M.J., Bienko, M., Dojer, N., Wang, Q., Karaca, E., Chiarle, R., Skrzypczak, M., Ginalski, K., et al. (2013). Nucleotide-resolution DNA double-strand break mapping by next-generation sequencing. Nat. Methods 10, 361–365.

Cullot, G., Boutin, J., Toutain, J., Prat, F., Pennamen, P., Rooryck, C., Teichmann, M., Rousseau, E., Lamrissi-Garcia, I., Guyonnet-Duperat, V., et al. (2019). CRISPR-Cas9 genome editing induces megabase-scale chromosomal truncations. Nat. Commun. 10, 1136.

Doench, J.G., Hartenian, E., Graham, D.B., Tothova, Z., Hegde, M., Smith, I., Sullender, M., Ebert, B.L., Xavier, R.J., and Root, D.E. (2014). Rational design of highly active sgRNAs for CRISPR-Cas9-mediated gene inactivation. Nat. Biotechnol. 32, 1262–1267.

Edraki, A., Mir, A., Ibraheim, R., Gainetdinov, I., Yoon, Y., Song, C.-Q., Cao, Y., Gallant, J., Xue, W., Rivera-Pérez, J.A., et al. (2018). A Compact, High-Accuracy Cas9 with a Dinucleotide PAM for In Vivo Genome Editing. Mol. Cell 73, 714–726.e4.

Fu, B.X.H., St. Onge, R.P., Fire, A.Z., and Smith, J.D. (2016). Distinct patterns of Cas9 mismatch tolerance in vitro and in vivo. Nucleic Acids Res. 44, 5365–5377.

Fu, Y., Foden, J.A., Khayter, C., Maeder, M.L., Reyon, D., Joung, J.K., and Sander, J.D. (2013). High-frequency off-target mutagenesis induced by CRISPR-Cas nucleases in human cells. Nat. Biotechnol. 31, 822–826.

Gao, P., Yang, H., Rajashankar, K.R., Huang, Z., and Patel, D.J. (2016). Type V CRISPR-Cas Cpf1 endonuclease employs a unique mechanism for crRNA-mediated target DNA recognition. Cell Res. 26, 901–913.

Gilbert, L.A., Horlbeck, M.A., Adamson, B., Villalta, J.E., Chen, Y., Whitehead, E.H., Guimaraes, C., Panning, B., Ploegh, H.L., Bassik, M.C., et al. (2014). Genome-Scale CRISPR-Mediated Control of Gene Repression and Activation. Cell 159, 647–661.

Gong, S., Yu, H.H., Johnson, K.A., and Taylor, D.W. (2018). DNA Unwinding Is the Primary Determinant of CRISPR-Cas9 Activity. Cell Rep. 22, 359–371.

Guenther, U.-P., Yandek, L.E., Niland, C.N., Campbell, F.E., Anderson, D., Anderson, V.E., Harris, M.E., and Jankowsky, E. (2013). Hidden specificity in an apparently nonspecific RNA-binding protein. Nature 502, 385–388.

Hawkins, J.A., Jones, S.K., Finkelstein, I.J., and Press, W.H. (2018). Indel-correcting DNA barcodes for high-throughput sequencing. Proc. Natl. Acad. Sci. 115, E6217–E6226.

Hsu, P.D., Scott, D.A., Weinstein, J.A., Ran, F.A., Konermann, S., Agarwala, V., Li, Y., Fine, E.J., Wu, X., Shalem, O., et al. (2013). DNA targeting specificity of RNA-guided Cas9 nucleases. Nat. Biotechnol. 31, 827–832.

Jiang, F., Taylor, D.W., Chen, J.S., Kornfeld, J.E., Zhou, K., Thompson, A.J., Nogales, E., and Doudna, J.A. (2016). Structures of a CRISPR-Cas9 R-loop complex primed for DNA cleavage. Science 351, 867–871.

Jiang, W., Bikard, D., Cox, D., Zhang, F., and Marraffini, L.A. (2013). RNA-guided editing of bacterial genomes using CRISPR-Cas systems. Nat. Biotechnol. 31, 233–239.

Jinek, M., Chylinski, K., Fonfara, I., Hauer, M., Doudna, J.A., and Charpentier, E. (2012). A Programmable Dual-RNA–Guided DNA Endonuclease in Adaptive Bacterial Immunity. Science 337, 816–821.

Jung, C., Hawkins, J.A., Jones, S.K., Xiao, Y., Rybarski, J.R., Dillard, K.E., Hussmann, J., Saifuddin, F.A., Savran, C.A., Ellington, A.D., et al. (2017). Massively Parallel Biophysical Analysis of CRISPR-Cas Complexes on Next Generation Sequencing Chips. Cell 170, 35–47.e13.

Kim, D., Bae, S., Park, J., Kim, E., Kim, S., Yu, H.R., Hwang, J., Kim, J.-I., and Kim, J.-S. (2015). Digenome-seq: genome-wide profiling of CRISPR-Cas9 off-target effects in human cells. Nat. Methods 12, 237–243.

Kim, D., Kim, J., Hur, J.K., Been, K.W., Yoon, S.-H., and Kim, J.-S. (2016). Genome-wide analysis reveals specificities of Cpf1 endonucleases in human cells. Nat. Biotechnol. 34, 863–868.

Kim, H.K., Song, M., Lee, J., Menon, A.V., Jung, S., Kang, Y.-M., Choi, J.W., Woo, E., Koh, H.C., Nam, J.-W., et al. (2017a). In vivo high-throughput profiling of CRISPR-Cpf1 activity. Nat. Methods 14, 153–159.

Kim, S., Bae, T., Hwang, J., and Kim, J.-S. (2017b). Rescue of high-specificity Cas9 variants using sgRNAs with matched 5’ nucleotides. Genome Biol. 18.

Kimsey, I.J., Petzold, K., Sathyamoorthy, B., Stein, Z.W., and Al-Hashimi, H.M. (2015). Visualizing transient Watson-Crick-like mispairs in DNA and RNA duplexes. Nature 519, 315–320.

Kleinstiver, B.P., Prew, M.S., Tsai, S.Q., Topkar, V.V., Nguyen, N.T., Zheng, Z., Gonzales, A.P.W., Li, Z., Peterson, R.T., Yeh, J.-R.J., et al. (2015). Engineered CRISPR-Cas9 nucleases with altered PAM specificities. Nature 523, 481–485.

Kleinstiver, B.P., Pattanayak, V., Prew, M.S., Tsai, S.Q., Nguyen, N.T., Zheng, Z., and Joung, J.K. (2016a). High-fidelity CRISPR-Cas9 nucleases with no detectable genome-wide off-target effects. Nature 529, 490–495.

Kleinstiver, B.P., Tsai, S.Q., Prew, M.S., Nguyen, N.T., Welch, M.M., Lopez, J.M., McCaw, Z.R., Aryee, M.J., and Joung, J.K. (2016b). Genome-wide specificities of CRISPR-Cas Cpf1 nucleases in human cells. Nat. Biotechnol. 34, 869–874.

Klompe, S.E., Vo, P.L.H., Halpin-Healy, T.S., and Sternberg, S.H. (2019). Transposon-encoded CRISPR–Cas systems direct RNA-guided DNA integration. Nature 1.

Komor, A.C., Kim, Y.B., Packer, M.S., Zuris, J.A., and Liu, D.R. (2016). Programmable editing of a target base in genomic DNA without double-stranded DNA cleavage. Nature advance online publication.

Lee, J.K., Jeong, E., Lee, J., Jung, M., Shin, E., Kim, Y., Lee, K., Jung, I., Kim, D., Kim, S., et al. (2018). Directed evolution of CRISPR-Cas9 to increase its specificity. Nat. Commun. 9, 3048.

Li, S.-Y., Cheng, Q.-X., Liu, J.-K., Nie, X.-Q., Zhao, G.-P., and Wang, J. (2018). CRISPR-Cas12a has both cis - and trans-cleavage activities on single-stranded DNA. Cell Res. 28, 491.

Lin, Y., Cradick, T.J., Brown, M.T., Deshmukh, H., Ranjan, P., Sarode, N., Wile, B.M., Vertino, P.M., Stewart, F.J., and Bao, G. (2014). CRISPR/Cas9 systems have off-target activity with insertions or deletions between target DNA and guide RNA sequences. Nucleic Acids Res. 42, 7473–7485.

Liu, M.-S., Gong, S., Yu, H.-H., Jung, K., Johnson, K.A., and Taylor, D.W. (2019). Basis for discrimination by engineered CRISPR/Cas9 enzymes. BioRxiv 630509.

Moreno-Mateos, M.A., Vejnar, C.E., Beaudoin, J.-D., Fernandez, J.P., Mis, E.K., Khokha, M.K., and Giraldez, A.J. (2015). CRISPRscan: designing highly efficient sgRNAs for CRISPR-Cas9 targeting in vivo. Nat. Methods 12, 982–988.

Murugan, K., Seetharam, A.S., Severin, A.J., and Sashital, D.G. (2019). Pervasive off-target and double-stranded DNA nicking by CRISPR-Cas12a. BioRxiv 657791.

Nishimasu, H., Ran, F.A., Hsu, P.D., Konermann, S., Shehata, S.I., Dohmae, N., Ishitani, R., Zhang, F., and Nureki, O. (2014). Crystal structure of Cas9 in complex with guide RNA and target DNA. Cell 156, 935–949.

Ozer, A., Tome, J.M., Friedman, R.C., Gheba, D., Schroth, G.P., and Lis, J.T. (2015). Quantitative Assessment of RNA-Protein Interactions with High Throughput Sequencing - RNA Affinity Profiling (HiTS-RAP). Nat. Protoc. 10, 1212–1233.

Pattanayak, V., Lin, S., Guilinger, J.P., Ma, E., Doudna, J.A., and Liu, D.R. (2013). High-throughput profiling of off-target DNA cleavage reveals RNA-programmed Cas9 nuclease specificity. Nat. Biotechnol. 31, 839–843.

Qi, L.S., Larson, M.H., Gilbert, L.A., Doudna, J.A., Weissman, J.S., Arkin, A.P., and Lim, W.A. (2013). Repurposing CRISPR as an RNA-Guided Platform for Sequence-Specific Control of Gene Expression. Cell 152, 1173–1183.

Ran, F.A., Cong, L., Yan, W.X., Scott, D.A., Gootenberg, J.S., Kriz, A.J., Zetsche, B., Shalem, O., Wu, X., Makarova, K.S., et al. (2015). In vivo genome editing using Staphylococcus aureus Cas9. Nature 520, 186–191.

Shmakov, S., Abudayyeh, O.O., Makarova, K.S., Wolf, Y.I., Gootenberg, J.S., Semenova, E., Minakhin, L., Joung, J., Konermann, S., Severinov, K., et al. (2015). Discovery and Functional Characterization of Diverse Class 2 CRISPR-Cas Systems. Mol. Cell 60, 385–397.

Slaymaker, I.M., Gao, L., Zetsche, B., Scott, D.A., Yan, W.X., and Zhang, F. (2016). Rationally engineered Cas9 nucleases with improved specificity. Science 351, 84–88.

Smargon, A.A., Cox, D.B.T., Pyzocha, N.K., Zheng, K., Slaymaker, I. M., Gootenberg, J.S., Abudayyeh, O.A., Essletzbichler, P., Shmakov, S., Makarova, K.S., et al. (2017). Cas13b Is a Type VI-B CRISPR-Associated RNA-Guided RNase Differentially Regulated by Accessory Proteins Csx27 and Csx28. Mol. Cell 65, 618–630.e7.

Stella, S., Mesa, P., Thomsen, J., Paul, B., Alcón, P., Jensen, S.B., Saligram, B., Moses, M.E., Hatzakis, N.S., and Montoya, G. (2018). Conformational Activation Promotes CRISPR-Cas12a Catalysis and Resetting of the Endonuclease Activity. Cell 175, 1856–1871.e21.

Sternberg, S.H., Redding, S., Jinek, M., Greene, E.C., and Doudna, J.A. (2014). DNA interrogation by the CRISPR RNA-guided endonuclease Cas9. Nature 507, 62–67.

Sternberg, S.H., LaFrance, B., Kaplan, M., and Doudna, J.A. (2015). Conformational control of DNA target cleavage by CRISPR-Cas9. Nature 527, 110–113.

Stormo, G.D., and Zhao, Y. (2010). Determining the specificity of protein-DNA interactions. Nat. Rev. Genet. 11, 751–760.

Strecker, J., Ladha, A., Gardner, Z., Schmid-Burgk, J.L., Makarova, K.S., Koonin, E.V., and Zhang, F. (2019). RNA-guided DNA insertion with CRISPR-associated transposases. Science eaax9181.

Strohkendl, I., Saifuddin, F.A., Rybarski, J.R., Finkelstein, I.J., and Russell, R. (2018). Kinetic Basis for DNA Target Specificity of CRISPR-Cas12a. Mol. Cell 71, 816–824.e3.

Sugimoto, N., Yasumatsu, I., and Fujimoto, M. (1997). Stabilities of internal rU-dG and rG-dT pairs in RNA/DNA hybrids. Nucleic Acids Symp. Ser. 199–200.

Sugimoto, N., Nakano, M., and Nakano, S. (2000). Thermodynamics-Structure Relationship of Single Mismatches in RNA/DNA Duplexes†. Biochemistry 39, 11270–11281.

Swarts, D.C., and Jinek, M. (2018). Mechanistic Insights into the cis-and trans-Acting DNase Activities of Cas12a. Mol. Cell 73, 589–600.e4.

Tsai, S.Q., Zheng, Z., Nguyen, N.T., Liebers, M., Topkar, V.V., Thapar, V., Wyvekens, N., Khayter, C., Iafrate, A.J., Le, L.P., et al. (2015). GUIDE-seq enables genome-wide profiling of off-target cleavage by CRISPR-Cas nucleases. Nat. Biotechnol. 33, 187–197.

Tsai, S.Q., Nguyen, N.T., Malagon-Lopez, J., Topkar, V.V., Aryee, M.J., and Joung, J.K. (2017). CIRCLE-seq: a highly sensitive *in vitro* screen for genome-wide CRISPR–Cas9 nuclease off-targets. Nat. Methods 14, 607–614.

Tycko, J., Myer, V.E., and Hsu, P.D. (2016). Methods for Optimizing CRISPR-Cas9 Genome Editing Specificity. Mol. Cell 63, 355–370.

Wang, T., Wei, J.J., Sabatini, D.M., and Lander, E.S. (2014). Genetic Screens in Human Cells Using the CRISPR-Cas9 System. Science 343, 80–84.

Wu, W.Y., Lebbink, J.H.G., Kanaar, R., Geijsen, N., and van der Oost, J. (2018). Genome editing by natural and engineered CRISPR-associated nucleases. Nat. Chem. Biol. 14, 642–651.

Xu, X., Duan, D., and Chen, S.-J. (2017). CRISPR-Cas9 cleavage efficiency correlates strongly with target-sgRNA folding stability: from physical mechanism to off-target assessment. Sci. Rep. 7, 143.

Yamano, T., Nishimasu, H., Zetsche, B., Hirano, H., Slaymaker, I.M., Li, Y., Fedorova, I., Nakane, T., Makarova, K.S., Koonin, E.V., et al. (2016). Crystal Structure of Cpf1 in Complex with Guide RNA and Target DNA. Cell 165, 949–962.

Yan, W.X., Mirzazadeh, R., Garnerone, S., Scott, D., Schneider, M.W., Kallas, T., Custodio, J., Wernersson, E., Li, Y., Gao, L., et al. (2017). BLISS is a versatile and quantitative method for genome-wide profiling of DNA double-strand breaks. Nat. Commun. 8, 15058.

Zeng, Y., Cui, Y., Zhang, Y., Zhang, Y., Liang, M., Chen, H., Lan, J., Song, G., and Lou, J. (2018). The initiation, propagation and dynamics of CRISPR-SpyCas9 R-loop complex. Nucleic Acids Res. 46, 350–361.

Zetsche, B., Gootenberg, J.S., Abudayyeh, O.O., Slaymaker, I.M., Makarova, K.S., Essletzbichler, P., Volz, S.E., Joung, J., van der Oost, J., Regev, A., et al. (2015). Cpf1 is a single RNA-guided endonuclease of a class 2 CRISPR-Cas system. Cell 163, 759–771.

Zhang, Y., Ge, X., Yang, F., Zhang, L., Zheng, J., Tan, X., Jin, Z.-B., Qu, J., and Gu, F. (2014). Comparison of non-canonical PAMs for CRISPR/Cas9-mediated DNA cleavage in human cells. Sci. Rep. 4, 5405.

