## Supplemental figures and text for "Massively parallel kinetic profiling of natural and engineered CRISPR nucleases"

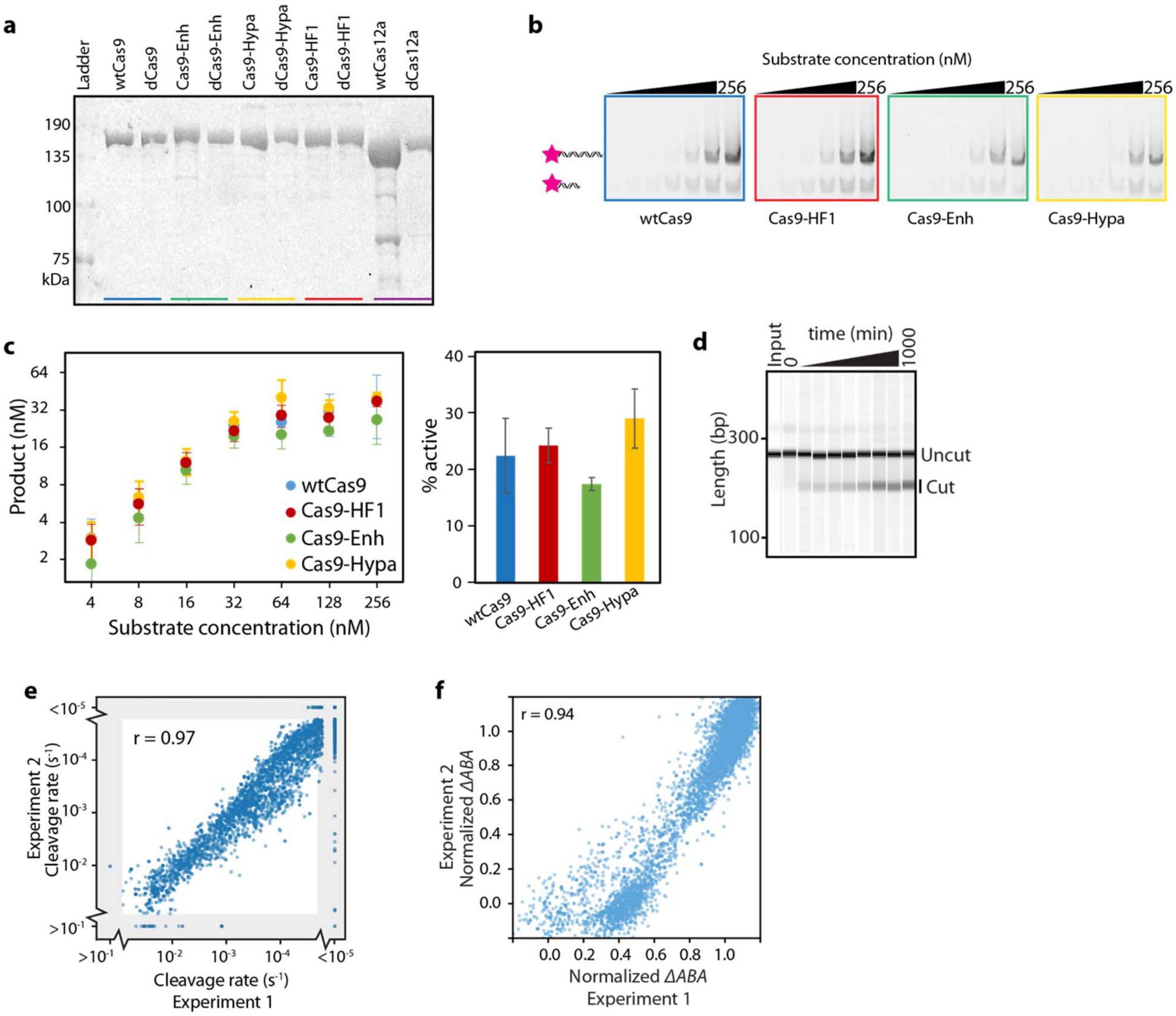


**Figure S1. Biochemical characterization of CRISPR-Cas nucleases. (a)** Coomassie-stained 10% SDS-PAGE gel of purified nucleases and their catalytically inactive variants. **(b)** Representative active site titration cleavage gels (10% TBE) and **(c)** quantification of three replicates for the indicated Cas9 variants. To determine the activity of each nuclease, 128 nM of the Cas9 RNP was incubated with 2-256nM of ATTO647N-labeled matched DNA (pink star) for 30 minutes. **(c, right)** The active nuclease concentration (mean ± SD; of at least three replicates) was determined from the concentration of product formed at 64-256 nM input DNA concentrations. **(d)** Time course of a wtCas9 nuclease reaction (sgRNA 2), as resolved by capillary electrophoresis. **(e)** Two independent wtCas12a NucleaSeq experiments show the same excellent reproducibility as for wtCas9 (see Figure 1G; Wald $\chi^2$ test statistic, excluding gray regions). Gray regions indicate sequences with cleavage rates that exceed the dynamic range of the experiment. **(f)** Two independent CHAMP experiments show excellent reproducibility (dCas9 with sgRNA 1, Wald $\chi^2$ test statistic).

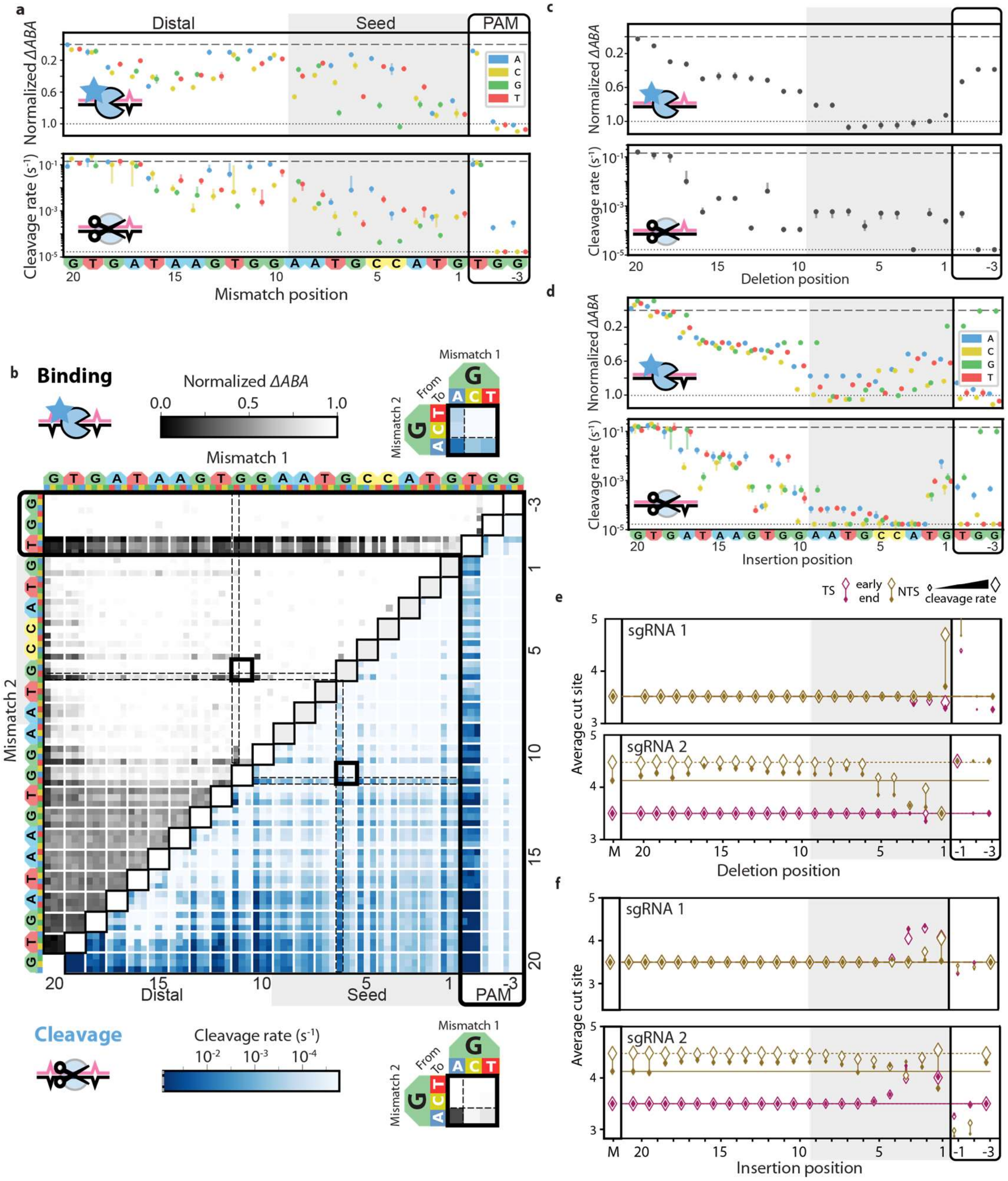
a
Distal
Seed
PAM
Normalized ΔABA
Cleavage rate (s⁻¹)
Mismatch position
A
C
G
T
b
Binding
Normalized ΔABA
0.0
0.5
1.0
Mismatch 1
Mismatch 2
From
To
Distal
Seed
PAM
Cleavage
Cleavage rate (s⁻¹)
c
Normalized ΔABA
Cleavage rate (s⁻¹)
Deletion position
d
Nnormalized ΔABA
Cleavage rate (s⁻¹)
Insertion position
e
TS
early end
NTS
cleavage rate
sgRNA 1
sgRNA 2
Average cut site
Deletion position
f
sgRNA 1
sgRNA 2
Average cut site
Insertion position

**Figure S2. Comprehensive analysis of off-target wtCas9 DNA binding and cleavage with sgRNA 2. (a)** dCas9 ΔABAs (upper, 0 indicates matched target) and cleavage rates (lower) for all DNAs with a single mismatch relative to sgRNA 2. Dotted line: normalized matched target ΔABA or cleavage rate. Dashed line: scrambled DNA ΔABA (negative control) or limit of detection for the slowest-cleaving targets. Error bars: SD (upper) or 5% to 95% confidence intervals (lower) as measured by bootstrap analysis. **(b)** ΔABAs (upper, grays) and cleavage rates (lower, blues) for DNAs containing two mismatches relative to sgRNA 2. Black boxes expanded in callouts. **(c)** dCas9 ΔABAs (upper) and Cas9 cleavage rates (lower) for DNAs containing a single nucleotide deletion or **(d)** a single nucleotide insertion compared to sgRNA 2. Error bars: ΔABA SD and cleavage rate 5% to 95% confidence intervals as measured by bootstrap analysis. **(e)** Average cut site positions for each strand (TS, NTS) from DNAs containing one deletion or **(f)** insertion compared to sgRNA 1 (upper) or 2 (lower). Range spans the first timepoint (early, open diamonds) to the final time point (end, filled diamonds). Diamond size indicates the average cleavage rates of the associated DNAs. Dashed and solid horizontal lines indicate average cut site positions for matched DNA at early and late time points.

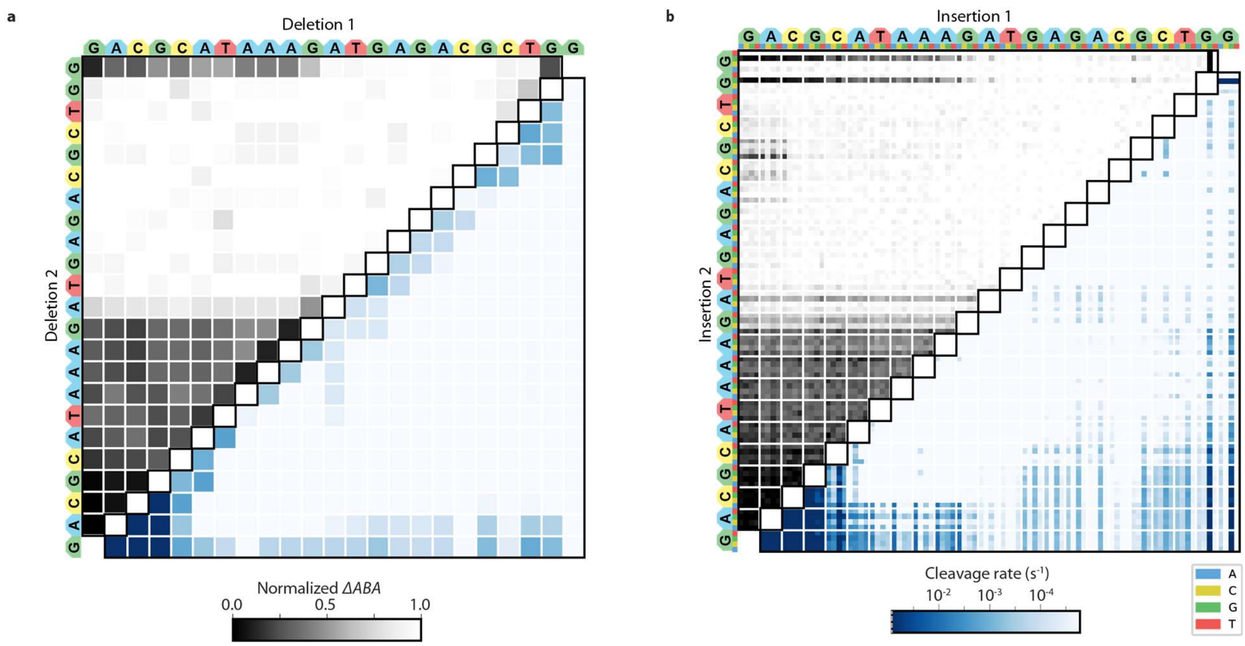


**Figure S3 Comprehensive analysis of off-target wtCas9 DNA binding and cleavage of DNAs with insertions or deletions. (a)** ΔABAs (upper) and cleavage rates (lower) for DNAs containing two deletions or **(b)** two insertions relative to sgRNA 1.

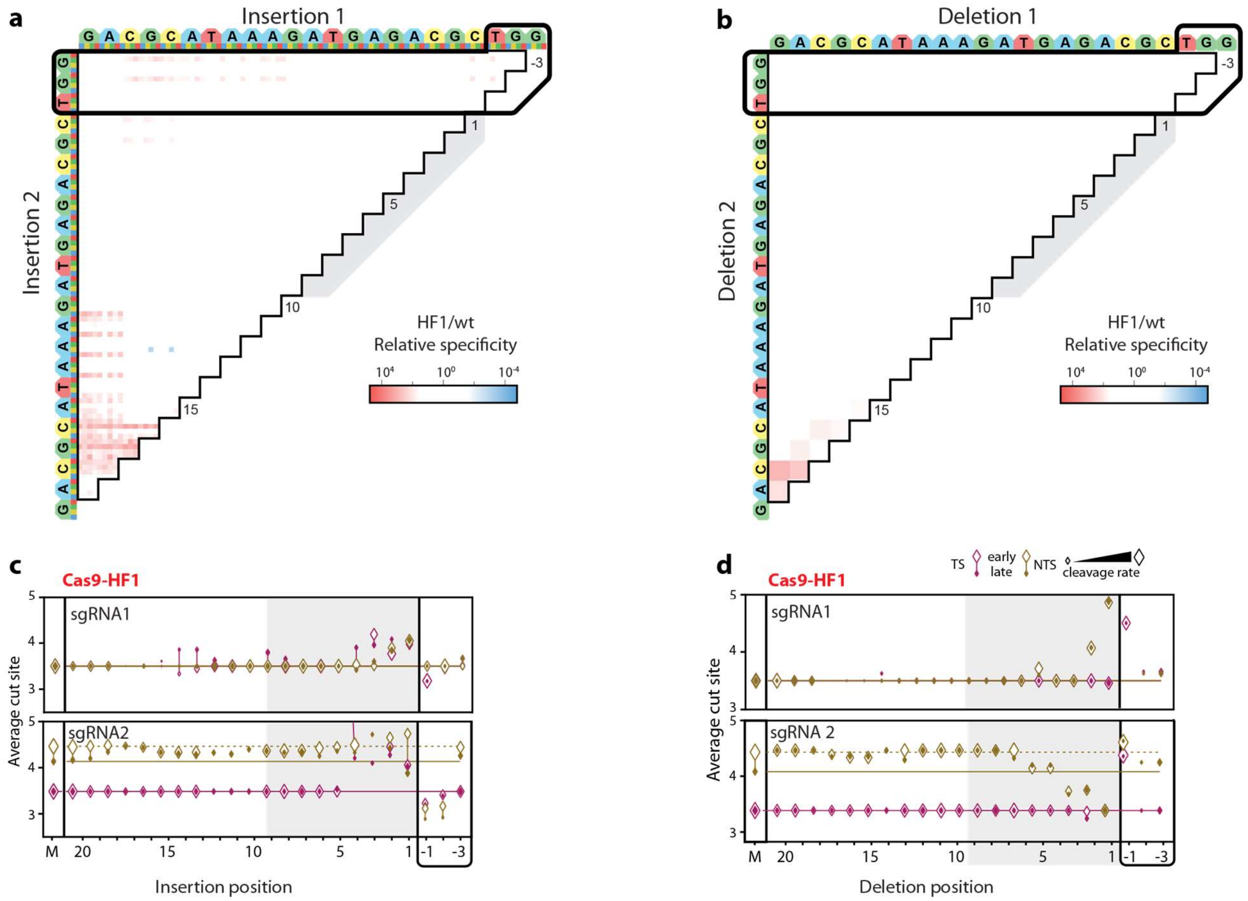


**Figure S4. Comparison of Cas9-HF1 and wtCas9 nuclease activities. (a)** Ratio of Cas9-HF1 to wtCas9 cleavage rates for all DNAs containing two insertions or **(b)** deletions compared to sgRNA 1. Red: slower cleavage by Cas9-HF1; blue: slower cleavage by wtCas9. **(c)** Average cut site positions generated by Cas9-HF1 for each strand (TS, NTS) for DNAs containing the insertions or **(d)** deletions compared to sgRNA 1 (upper) or sgRNA 2 (lower). Range spans the first timepoint (early, open diamonds) to the final time point (late, filled diamonds). Diamond size indicates the average cleavage rates of the associated DNAs. Dashed and solid horizontal lines indicate average cut site positions for matched DNA at early and late time points.

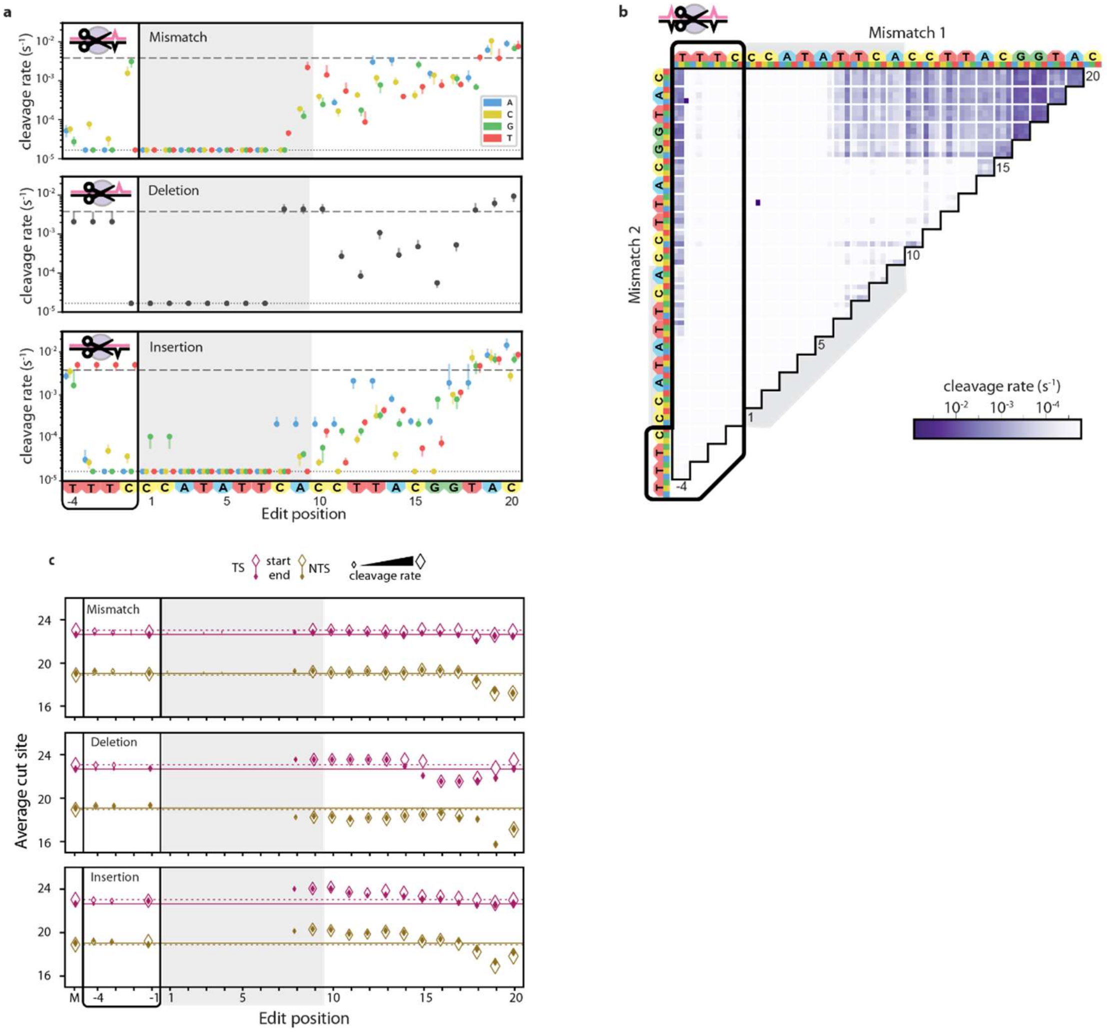


**Figure S5. Analysis of off-target wtCas12a cleavage with crRNA 4.**

**(a)** Cleavage rates for all DNAs with a single mismatch (upper), deletion (middle) or insertion (lower). Dotted line: matched target cleavage rate. Dashed line: limit of detection for the slowest-cleaving targets. Error bars: 5% to 95% confidence intervals as measured by bootstrap analysis. **(b)** Cleavage rates for DNAs containing two mismatches. **(c)** Average cut site positions generated by wtCas12a for each strand (TS, NTS) for DNAs containing the mismatches (upper), deletions (middle) or insertions (lower). Range spans the first timepoint (early, open diamonds) to the final time point (late, filled diamonds). Diamond size indicates the average cleavage rates of the associated DNAs, as in (a). Dashed and solid horizontal lines indicate average cut site positions for matched DNA at early and late time points.

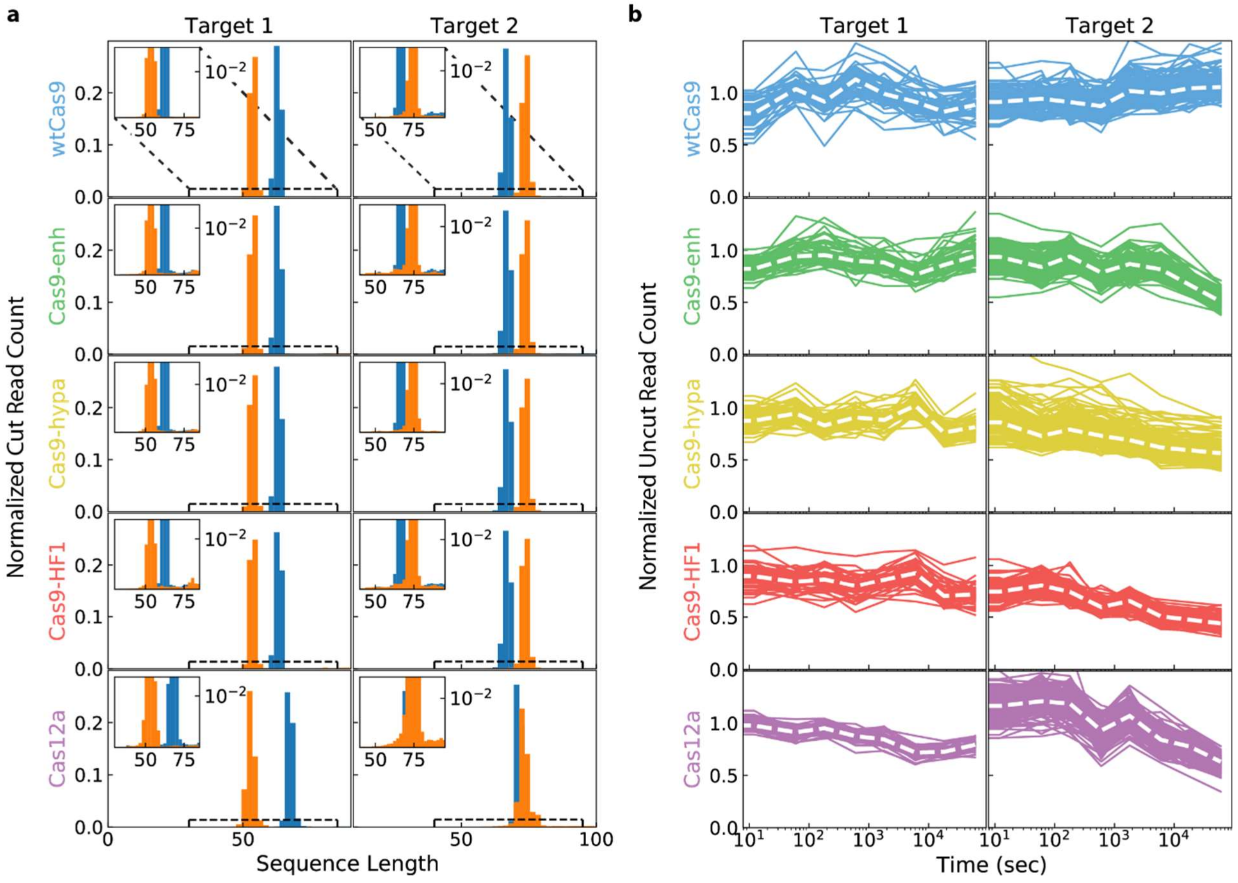


**Figure S6. Cas12a exhibits limited *trans* cleavage of NucleaSeq libraries.** We looked for two signatures of *trans* cleavage activity. (a) First, the distribution of normalized read counts of DNAs that only have the left (blue) or right (orange) barcodes show limited cut fragments outside the expected enzyme cleavage sites. Inset: a zoomed in view shows that there are few indiscriminately cut products outside of the main peaks, which correspond to the canonical cut site. A comparable amount of short DNA for Cas9 and Cas12 suggests that these fragments arise during library preparation and NGS. Cas12a-catalyzed *trans* cleaved DNA is a relatively minor component of our data (within the noise). Histograms are normalized to have a total area of one. (b) Second, we looked at the time-dependent read counts of ~150 uncut control sequences that are not complementary to the sg/crRNAs (overlapping lines). These sequences are normalized only for total read counts at each time, then with the value at time zero set to one. These values are proportional to the fraction of the library at each time point, not absolute read count. Robust *trans* cleavage by Cas12a would be expected to deplete these values over time at a faster rate than other proteins. However, Cas12a behaves similarly to the engineered Cas9 variants in all cases. Furthermore, we do not observe any individual traces decreasing faster than the group, as would be observed if trans-cleavage showed sequence bias for some of the control sequences. White dashed line: median.

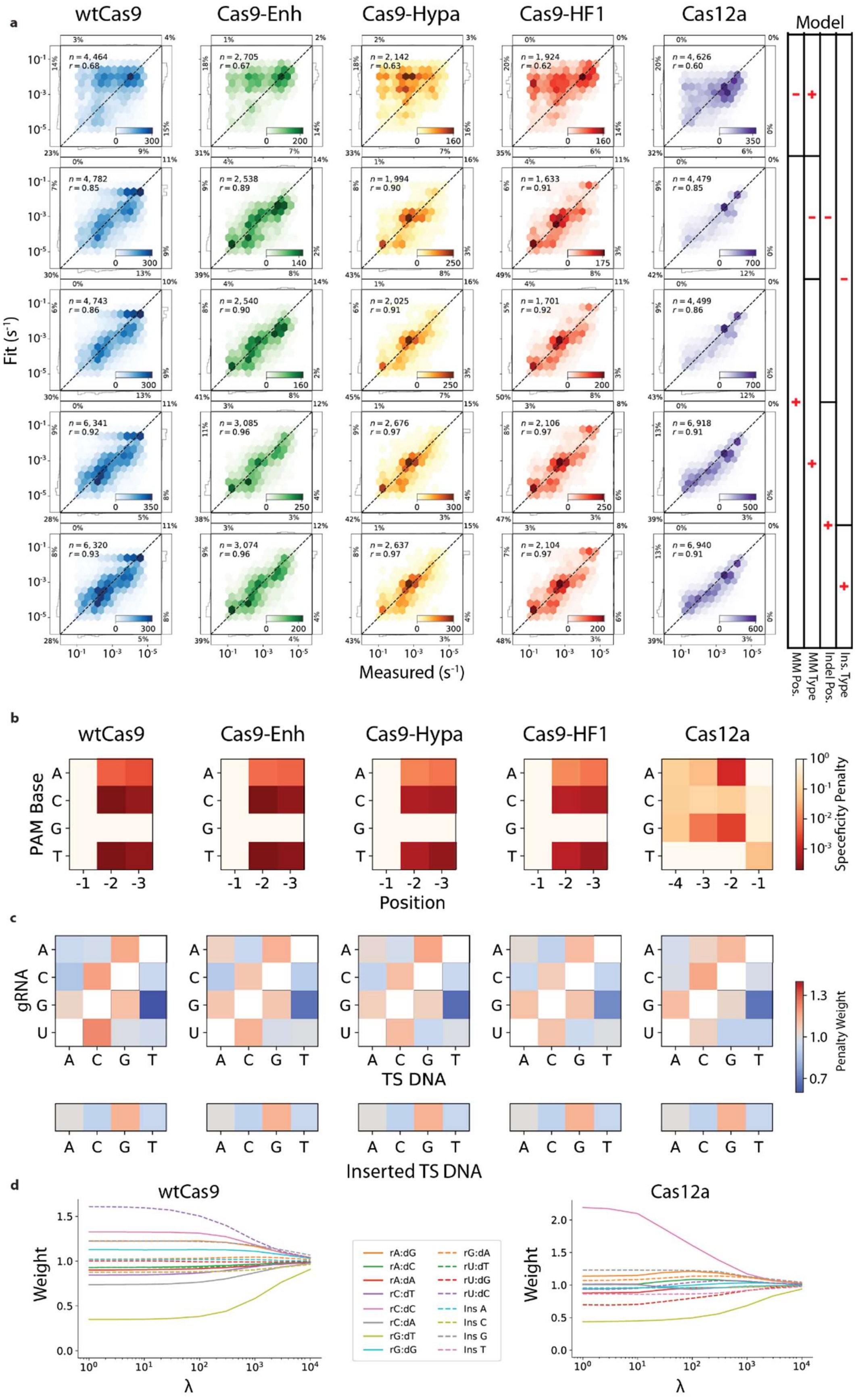
a
wtCas9
Cas9-Enh
Cas9-Hypa
Cas9-HF1
Cas12a
Model
Fit (s-1)
Measured (s-1)
MM Pos.
MM Type
Indel Pos.
Ins. Type
b
PAM Base
Position
Specificity Penalty
c
gRNA
TS DNA
Penalty Weight
Inserted TS DNA
d
Weight
λ
rA:dG
rA:dC
rA:dA
rC:dT
rC:dC
rC:dA
rG:dT
rG:dG
rG:dA
rU:dT
rU:dG
rU:dC
Ins A
Ins C
Ins G
Ins T

**Figure S7. Comparison of biophysical models for Cas nuclease specificity. (a)** Correlation between measured and fit cleavage rates for each protein for the indicated simplified model. (**b**) PAM position weight matrices computed from the cleavage rates for each protein. (**c**) Penalty weight values of mismatch (top) and insertion types (bottom) for each protein. (**d**) Transition and insertion weights as a function of the regularization parameter, $\lambda$ (see supplemental computational methods).

$$\frac{|C|_t^{side}}{Z_t^{side}} = k^{side}\left(1 - \frac{|F|_t}{|F|_0}\right)$$

We choose to set the final normalization constant $Z_{t_f}^{side} = 1$ and solve the above for $k^{side}$. Plugging this back in and rearranging gives normalization constants

$$Z_t^{side} = \frac{|C|_t^{side}}{|C|_{t_f}^{side}}\left(\frac{1 - |F|_{t_f}/|F|_0}{1 - |F|_t/|F|_0}\right)$$

This is intentionally a function only of ratios of read counts, not absolute read counts. This lets us use the median read count ratios from all 146 matched target controls to calculate the normalization constants. These final normalization constants are then used for all library members. Finally, read counts are normalized to range between zero and one. For full-length products, we normalize by the fit value of reads at time zero. For cleaved products, we normalize first by the sum of all cleaved products at all time points, then normalize to set the resulting median sum of all cleaved products at the final time point to the depletion of full-length products, $1 - |F|_{t_f}/|F|_0$.

The normalized read counts were fit to a single exponential decay. We observed that the data was well described by a single exponential, implying a constant reaction rate under the single turnover conditions used in this assay. A small fraction of the starting DNA sequences of each species were never cleaved, possibly indicating some hydrolytically inactive enzymes. We thus fit for exponential decay with a constant offset. For the constant offset, we used the median normalized fraction of uncleaved sequences of the 146 perfect target sequences at the final time point. Error bars give the standard deviation of 50 bootstrap measurements, each of which was calculated by resampling the raw read counts with replacement, renormalizing, and refitting (Efron and Tibshirani, 1993).

**Modeling cleavage specificity**

*Model description*

We modeled cleavage specificity (*Model 1*), given as the ratio of the cleavage rate of a given sequence $s$, $k_s$, to the cleavage rate of the matched sequence $m$, $k_m$, as:

$$\log\frac{k_s}{k_m} = \sum_{i\in\mathcal{P}} \log\Lambda(i, s_i) + \sum_{i\in\mathcal{D}} \log P_D(i) + \sum_{i\in\mathcal{I}} w_I(s_i)\log P_I(i) + \sum_{i\in\mathcal{M}} t_M(r_i, s_i)\log P_M(i)$$

The terms of the model give cleavage rate penalties for the following sequence edits respectively: suboptimal bases in the PAM, target deletions, target insertions, and target mismatches, each with corresponding set of positions with the given sequence edit type: $\mathcal{P}$, $\mathcal{D}, \mathcal{J}$, and $\mathcal{M}$. For suboptimal PAM bases, the cleavage rate penalty is given by the function $\Lambda$, a function of both the suboptimal base identity, $s_i$, and its position $i$.

$$\begin{bmatrix} n \\ - \\ x \end{bmatrix} = H_n w, \quad w = \frac{1}{n} H_n^T \begin{bmatrix} n \\ - \\ x \end{bmatrix}$$

Cleavage rates that are shorter than the first time point or longer than the last one cannot be modeled accurately. We therefore constrained the output of our models with the following "bandpass filter" function:

$$B(x) = \begin{cases} x & s \leq x \leq f \\ s & x < s \\ f & x > f \end{cases}$$

Where $s$ and $f$ are the slowest and fastest detectible cleavage rates, corresponding to half-lives at our first and last time points.

Ridge regularization of the difference of insertion and mismatch weights from one was used to reduce over-fitting of the underlying cleavage data (Hoerl and Kennard, 1970). Figure S7D shows the fit weight values as a function of the regularization parameter $\lambda$. The relative parameter values appear to stabilize near $\lambda = 10^3$, which we used to fit the model.

*Simplified models*

For comparison, we fit our data to four simplified models, each excluding some terms and/or factors in the full model above. The first three simplified models did not include the insertion or deletion terms, modeling the possibility that the recognition channel does not accommodate bulges to realign matching sequences after indels. Under this assumption, for example, a sequence with a single insertion between the first and second bases, but otherwise perfectly matching, would result in about 75% mismatches due to a forced frame shift. These three models were: cleavage rate as a function of only the mismatch base pair identities, only the mismatch position, or both as in the full model above. The fourth simplified model included insertions and deletions but omitted the insertion weights $w_I$. Each simplified model included the PAM term. We number the models for reference:

$$Model\ 5\colon\ \log \frac{k_s}{k_m} = \sum_{i \in \mathcal{P}} \log \Lambda(i, s_i) + \sum_{i \in \mathcal{M}} \log T_M(r_i, s_i)$$

$$Model\ 4\colon\ \log \frac{k_s}{k_m} = \sum_{i \in \mathcal{P}} \log \Lambda(i, s_i) + \sum_{i \in \mathcal{M}} \log P_M(i)$$

$$Model\ 3\colon\ \log \frac{k_s}{k_m} = \sum_{i \in \mathcal{P}} \log \Lambda(i, s_i) + \sum_{i \in \mathcal{M}} t_M(r_i, s_i) \log P_M(i)$$

$$Model\ 2\colon \log\frac{k_s}{k_m} = \sum_{i\in\mathcal{P}} \log\Lambda(i,s_i) + \sum_{i\in\mathcal{D}} \log P_D(i) + \sum_{i\in\mathcal{I}} \log P_I(i) + \sum_{i\in\mathcal{M}} t_M(r_i,s_i)\log P_M(i)$$

And Model 1 is the full model above. The mismatching base pairs function in Model 5, $T_M(r_i, s_i)$, is different from the analogous weighting function $t_M(r_i, s_i)$ in the other models as it gives absolute penalty values, not weights constrained to average value of one.

**Table S1. Nucleic acids used in this work**

| Name | RNP | Use | Sequence | References |
|---|---|---|---|---|
| sgRNA 1 | Cas9 | guide | UAAUACGACUCACUAUAGGACGCAUAAAGAUGAGACGCGUUUUAGAGCUAGAAAUAGCAAGUUAAAAUAAGGCUAGUCCGUUAUCAACUUGAAAAAGUGGCACCGAGUCGGUGCUUUU | This work |
| sgRNA 2 | Cas9 | guide | UAAUACGACUCACUAUAGGUGAUAAGUGGAAUGCCAUGGUUUUAGAGCUAGAAAUAGCAAGUUAAAAUAAGGCUAGUCCGUUAUCAACUUGAAAAAGUGGCACCGAGUCGGUGCUUUU | |
| crRNA 3 | Cas12a | guide | GUCAAAAGACCUUUUUAAUUUCUACUCUUGUAGAUGUGAUAAGUGGAAUGCCAUGUGGA | |
| crRNA 4 | Cas12a | guide | GUCAAAAGACCUUUUUAAUUUCUACUCUUGUAGAUGACGCAUAAAGAUGAGACGCUGGA | |
| pr238 | dCas9 | H840A | TCGTTTAAGTGATTATGATGTCGATGCCATTGTTCCACAAAGTTTCCTTAAA | (Jinek et al., 2012) |
| pr239 | | | TTTAAGGAAACTTTGTGGAACAATGGCATCGACATCATAATCACTTAAACGA | |
| pr236 | | D10A | GAAATACTCAATAGGCTTAGCTATCGGCACAAATAGCGTCG | |
| pr237 | | | CGACGCTATTTGTGCCGATAGCTAAGCCTATTGAGTATTTC | |
| pr271 | Cas9-enh | K848A | CGATCACATTGTTCCACAAAGTTTCCTTGCAGACGATTCAATAGACAATAAG | (Slaymaker et al., 2016) |
| pr272 | | | CTTATTGTCTATTGAATCGTCTGCAAGGAAACTTTGTGGAACAATGTGATCG | |

| | | | | |
|---|---|---|---|---|
| pr273 | | K1003A | CGTTGGAACTGCTTTGATTAAGAAATATCCAGCACTTGAATCGGAGTTTGT | |
| pr274 | | | ACAAACTCCGATTCAAGTGCTGGATATTTCTTAATCAAAGCAGTTCCAACG | |
| pr275 | | K1060A | ACTTGCAAATGGAGAGATTCGCAAAGCCCCTCTAATCGAA | |
| pr276 | | | TTCGATTAGAGGGGCTTTGCGAATCTCTCCATTTGCAAGT | |
| pr263 | Cas9-HF1 | N497A | CAGCTCAATCATTTATTGAACGCATGACAGCCTTTGATAAAAATCTTCCAAATGAAAAAG | (Kleinstiver et al., 2016a) |
| pr264 | | | CTTTTTCATTTGGAAGATTTTTATCAAAGGCTGTCATGCGTTCAATAAATGATTGAGCTG | |
| pr269 | | Q926A | CCAATTGGTTGAAACTCGCGCAATCACTAAGCATGTGGCA | |
| pr270 | | | TGCCACATGCTTAGTGATTGCGCGAGTTTCAACCAATTGG | |
| pr265 | | R661A | GCCGTTATACTGGTTGGGGAGCTTTGTCTCGAAAATTGATTA | |
| pr266 | | | TAATCAATTTTCGAGACAAAGCTCCCCAACCAGTATAACGGC | |
| pr267 | Cas9-HF1 | Q695A | GTTTTGCCAATCGCAATTTTATGGCGCTGATCCATGATGATAGTTTG | (Kleinstiver et al., 2016a) |
| pr268 | | | CAAACTATCATCATGGATCAGCGCCATAAAATTGCGATTGGCAAAAC | |
| pr399 | Cas9-hypa | H698A, Q695A | GCGCTGATCGCTGATGATAGTTTGACATTTAAAGAAGACATTCAAAAAGCACAA | (Chen et al., 2017) |

| | | | | |
|---|---|---|---|---|
| pr400 | | M694A, N692A | GCCAAAGCTGCGATTGGCAAAACCA TCTGATTTCAAAAA | |
| pr309 | sgRNA 1, crRNA 4 | Matched target 1 | ACGCTCTTCCGATCTTTTAGACGCAT AAAGATGAGACGCTGGAGATCGGAA GAGCAC | This work |
| pr310 | | | GTGCTCTTCCGATCTCCAGCGTCTCA TCTTTATGCGTCTAAAAGATCGGAAG AGCGT | |
| pr477 | | Library 1 | Oligonucleotide pool. See table S3. | |
| pr475 | | Primer for library 1 | ATAACTAATTGAGCTGAACGCAC | |
| pr476 | | | CTGAATAGTCGTGTAGTTGTGCT | |
| pr307 | sgRNA 2, crRNA 3 | Matched target 2 | ACGCTCTTCCGATCTTTTAGTGATAA GTGGAATGCCATGTGGAGATCGGAA GAGCAC | |
| pr308 | | | GTGCTCTTCCGATCTCCACATGGCAT TCCACTTATCACTAAAAGATCGGAAG AGCGT | |
| pr371 | | Library 2 | Oligonucleotide pool. See table S3. | |
| pr364 | | Primer for library 2 | AACCGCCGAATAACAGAGT | |
| pr365 | | | AAGAACGCCTCGCACACT | |
| pr460 | | Atto647N Primer for library 2 | /5atto647n/AACCGCCGAATAACAGAGT | |

**Table S2. Plasmids used in this work.**

| Name | Protein | construct | Mutations | References |
|---|---|---|---|---|
| pIF324 | *Sp*Cas9 | 6xHis-MBP-3xFlag-*Sp*Cas9 | | (Jinek et al., 2012) |
| pIF335 | | 6xHis-MBP-3xFlag-d*Sp*Cas9 | D10A, H840A | (Jinek et al., 2012) |
| pIF325 | | 6xHis-MBP-3xFlag-enhanced *Sp*Cas9-1.1 | K848A, K1003A, R1060A | (Slaymaker et al., 2016) |
| pIF326 | | 6xHis-MBP-3xFlag-enhanced d*Sp*Cas9-1.1 | D10A, H840A, K848A, K1003A, R1060A | (Slaymaker et al., 2016) |
| pIF329 | | 6xHis-MBP-3xFlag-*Sp*Cas9-HF1 | R661A, Q695A, Q926A | (Kleinstiver et al., 2016a) |
| pIF330 | | 6xHis-MBP-3xFlag-d*Sp*Cas9-HF1 | D10A, R661A, Q695A, H840A, Q926A | (Kleinstiver et al., 2016a) |
| pIF350 | | 6xHis-MBP-3xFlag-hypa *Sp*Cas9 | N692A, M694A, Q695A, H698A | (Chen et al., 2017) |
| pIF351 | | 6xHis-MBP-3xFlag-hypa d*Sp*Cas9 | D10A, N692A, M694A, Q695A, H698A, H840A | (Chen et al., 2017) |
| pIF502 | *As*Cas12a | 6xHis-TwinStrep-SUMO-*As*Cas12a-3xFlag | | (Strohkendl et al., 2018) |

### REFERENCES


Akaike, H. (1974). A new look at the statistical model identification. IEEE Trans. Autom. Control *19*, 716–723.

Chen, J.S., Dagdas, Y.S., Kleinstiver, B.P., Welch, M.M., Sousa, A.A., Harrington, L.B., Sternberg, S.H., Joung, J.K., Yildiz, A., and Doudna, J.A. (2017). Enhanced proofreading governs CRISPR–Cas9 targeting accuracy. Nature *550*, 407–410.

Cieślik, M., Pederson, B., and Arindrarto, W. (2016). Align: polite, proper sequence alignment.

Cock, P.J.A., Antao, T., Chang, J.T., Chapman, B.A., Cox, C.J., Dalke, A., Friedberg, I., Hamelryck, T., Kauff, F., Wilczynski, B., et al. (2009). Biopython: freely available Python tools for computational molecular biology and bioinformatics. Bioinformatics *25*, 1422–1423.

Edelstein, A.D., Tsuchida, M.A., Amodaj, N., Pinkard, H., Vale, R.D., and Stuurman, N. (2014). Advanced methods of microscope control using μManager software. J. Biol. Methods *1*, 10.

Efron, B., and Tibshirani, R.J. (1993). An Introduction to the Bootstrap (New York: Chapman and Hall/CRC).

Fu, B.X.H., St. Onge, R.P., Fire, A.Z., and Smith, J.D. (2016). Distinct patterns of Cas9 mismatch tolerance in vitro and in vivo. Nucleic Acids Res. *44*, 5365–5377.

Hawkins, J.A., Jones, S.K., Finkelstein, I.J., and Press, W.H. (2018). Indel-correcting DNA barcodes for high-throughput sequencing. Proc. Natl. Acad. Sci. *115*, E6217–E6226.

Hoerl, A.E., and Kennard, R.W. (1970). Ridge Regression: Biased Estimation for Nonorthogonal Problems. Technometrics *12*, 55–67.

Hsu, P.D., Scott, D.A., Weinstein, J.A., Ran, F.A., Konermann, S., Agarwala, V., Li, Y., Fine, E.J., Wu, X., Shalem, O., et al. (2013). DNA targeting specificity of RNA-guided Cas9 nucleases. Nat. Biotechnol. *31*, 827–832.

Jinek, M., Chylinski, K., Fonfara, I., Hauer, M., Doudna, J.A., and Charpentier, E. (2012). A Programmable Dual-RNA–Guided DNA Endonuclease in Adaptive Bacterial Immunity. Science *337*, 816–821.

Jung, C., Hawkins, J.A., Jones, S.K., Xiao, Y., Rybarski, J.R., Dillard, K.E., Hussmann, J., Saifuddin, F.A., Savran, C.A., Ellington, A.D., et al. (2017). Massively Parallel Biophysical Analysis of CRISPR-Cas Complexes on Next Generation Sequencing Chips. Cell *170*, 35-47.e13.

Kim, D., Bae, S., Park, J., Kim, E., Kim, S., Yu, H.R., Hwang, J., Kim, J.-I., and Kim, J.-S. (2015). Digenome-seq: genome-wide profiling of CRISPR-Cas9 off-target effects in human cells. Nat. Methods *12*, 237–243.

Kim, D., Kim, J., Hur, J.K., Been, K.W., Yoon, S.-H., and Kim, J.-S. (2016). Genome-wide analysis reveals specificities of Cpf1 endonucleases in human cells. Nat. Biotechnol. *34*, 863–868.

Kim, H.K., Song, M., Lee, J., Menon, A.V., Jung, S., Kang, Y.-M., Choi, J.W., Woo, E., Koh, H.C., Nam, J.-W., et al. (2017). In vivo high-throughput profiling of CRISPR-Cpf1 activity. Nat. Methods *14*, 153–159.

Kleinstiver, B.P., Pattanayak, V., Prew, M.S., Tsai, S.Q., Nguyen, N.T., Zheng, Z., and Joung, J.K. (2016a). High-fidelity CRISPR-Cas9 nucleases with no detectable genome-wide off-target effects. Nature *529*, 490–495.

Kleinstiver, B.P., Tsai, S.Q., Prew, M.S., Nguyen, N.T., Welch, M.M., Lopez, J.M., McCaw, Z.R., Aryee, M.J., and Joung, J.K. (2016b). Genome-wide specificities of CRISPR-Cas Cpf1 nucleases in human cells. Nat. Biotechnol. *34*, 869–874.

Pattanayak, V., Lin, S., Guilinger, J.P., Ma, E., Doudna, J.A., and Liu, D.R. (2013). High-throughput profiling of off-target DNA cleavage reveals RNA-programmed Cas9 nuclease specificity. Nat. Biotechnol. *31*, 839–843.

Rohatgi, A. (2019). WebPlotDigitizer.

Schindelin, J., Arganda-Carreras, I., Frise, E., Kaynig, V., Longair, M., Pietzsch, T., Preibisch, S., Rueden, C., Saalfeld, S., Schmid, B., et al. (2012). Fiji: an open-source platform for biological-image analysis. Nat. Methods *9*, 676–682.

Slaymaker, I.M., Gao, L., Zetsche, B., Scott, D.A., Yan, W.X., and Zhang, F. (2016). Rationally engineered Cas9 nucleases with improved specificity. Science *351*, 84–88.

Strohkendl, I., Saifuddin, F.A., Rybarski, J.R., Finkelstein, I.J., and Russell, R. (2018). Kinetic Basis for DNA Target Specificity of CRISPR-Cas12a. Mol. Cell *71*, 816-824.e3.

Yan, W.X., Mirzazadeh, R., Garnerone, S., Scott, D., Schneider, M.W., Kallas, T., Custodio, J., Wernersson, E., Li, Y., Gao, L., et al. (2017). BLISS is a versatile and quantitative method for genome-wide profiling of DNA double-strand breaks. Nat. Commun. *8*, 15058.
